# Streptococcal natural products mediate interspecies competition with Gram-positive bacterial pathogens

**DOI:** 10.1101/2025.07.31.667707

**Authors:** Ruth Y. Isenberg, Celine W. Sackih, Julia L. E. Willett

## Abstract

Polymicrobial communities where bacteria must compete with each other to persist can serve as a source of uncharacterized antibacterial compounds to develop drugs for the treatment of drug-resistant infections. This study investigates interbacterial competition between bacteria found in the oral cavity, where *Streptococcus* species comprise a large portion of the resident oral microbiota and *Enterococcus faecalis* is a pathobiont that is commonly found in root canal infections. We used co-cultures to determine whether oral Streptococci and *E. faecalis* compete with each other. Our experiments revealed that multiple strains of *Streptococcus mutans*, an important cariogenic bacterium, kill vancomycin-resistant *Enterococcus* and other Gram-positive bacteria, including *Staphylococcus epidermidis*, and methicillin-resistant *Staphylococcus aureus*. Further, inhibition of Gram-positive bacteria by some strains of *S. mutans* requires the production of the non-ribosomal cyclic lipopeptide mutanobactin, while another strain inhibits independent of mutanobactin. We determined that *S. mutans* mutanobactin production increases target cell membrane permeability and that killing is contact-dependent. We also determined that an *E. faecalis* virulence factor, the secreted protease gelatinase (GelE), is required for recovery from mutanobactin-mediated killing. Additionally, data show that *S. mutans* mutanobactin production prevents and kills *E. faecalis* biofilms. Together, this work demonstrates how natural products from a common oral bacterium contribute to competition in polymicrobial environments, which will inform future strategies to treat and prevent bacterial infections.

**IMPORTANCE:** Antimicrobial resistance requires new therapeutics to treat drug-resistant infections. Novel antimicrobial compounds can be discovered in polymicrobial communities, where bacterial natural products promote competitive fitness. The oral cavity hosts a microbial consortium, and we investigated interactions between oral streptococci and *Enterococcus faecalis*, a common root canal infection isolate. We demonstrate antibacterial activity of streptococcal mutanobactin against Gram-positive pathogens, including antibiotic-resistant isolates. We further show that the *E. faecalis* virulence factor gelatinase promotes recovery from mutanobactin-mediated killing, and that mutanobactin prevents and kills *E. faecalis* biofilms. By probing interactions between bacteria that occupy the same niche and characterizing antibacterial activity of a bacterial product, this work contributes to broader efforts to identify and develop antibiotics to treat clinically relevant drug-resistant infections.

## INTRODUCTION

Multi-drug-resistant bacterial infections pose a significant threat to public health and place a large financial burden on healthcare systems. The ESKAPE pathogens (*Enterococcus faecium*, *Staphylococcus aureus*, *Klebsiella pneumoniae*, *Acinetobacter baumannii*, *Pseudomonas aeruginosa*, and *Enterobacter* spp.) are responsible for a large portion of hospital-acquired antibiotic resistant infections (1, 2). Additionally, of the 18 most alarming antibiotic resistance threats, the United States Centers for Disease Control and Prevention has identified six that cost the U.S. over 4.6 billion dollars per year (3). One of these threats is vancomycin-resistant *Enterococcus* (VRE). *Enterococcus faecalis* and *Enterococcus faecium* are Gram-positive gastrointestinal pathobionts that are leading causes of many hospital-acquired infections, such as endocarditis, bacteriemia, urinary tract infections, and wound/surgical site infections (4–12). VRE isolated from patients harbor a number of genetic elements that promote virulence, including the enterococcal pathogenicity island (13–19), and resistance to last-resort antibiotics such as vancomycin can lead to drug-resistant strain dominance in the gastrointestinal tract that can promote bloodstream infection following antibiotic treatment (12, 20–23). While antibiotics such as linezolid, daptomycin, and tigecycline are used to treat VRE, resistance to these drugs is of increasing concern (5, 19, 21, 23). Therefore, the identification and development of new drugs to combat antimicrobial resistance are of critical importance.

While *E. faecalis* is a gastrointestinal commensal bacterium, it is also found in the oral cavity and is frequently detected in the saliva and root canals of patients with periodontitis (24–39). Further, many oral isolates of *E. faecalis* are resistant to antibiotics, suggesting the oral cavity could act as a reservoir for virulent and drug-resistant *E. faecalis* that cause hospital-acquired infections (35). Understanding *E. faecalis* mechanisms for survival within the oral cavity will broaden our knowledge of *E. faecalis* strategies for persistence and subsequent dissemination to infection sites.

The oral cavity is host to a diverse consortium of species, with the *Streptococcus* genus among the most abundant. Some *Streptococcus* species colonize specific and distinct oral sites, while others are found in all oral sites but reach higher abundance in specific sites (40, 41). Additionally, *S. mutans* has been detected within infected root canals (42, 43). Because of the abundance and distribution of *Streptococcus* species within the oral cavity, the presence of *E. faecalis* in the oral cavity, and the presence of *E. faecalis* and *S. mutans* in infected root canals, we asked whether *Streptococcus* species compete with *E. faecalis* in co-culture. We determined that production of the non-ribosomal cyclic lipopeptide mutanobactin by *Streptococcus mutans* has antibacterial activity against *E. faecalis.* Subsequent co-culture experiments revealed that *S. mutans* antibacterial activity extends to all *Enterococcus* spp. and all Gram-positive species tested, including VRE, methicillin-resistant *S. aureus* (MRSA), and *Staphylococcus epidermidis*. We also demonstrate that mutanobactin-mediated killing prevents and reduces *E. faecalis* biofilms. All together, our work characterizes interbacterial competition between *E. faecalis* and *S. mutans* as well as describes a novel antibacterial function for the previously identified natural product mutanobactin. Our findings highlight the utility in examining well-characterized microbial consortia for undiscovered mechanisms of interbacterial competition mediated by natural products. These compounds may be leveraged into new treatments for antibiotic-resistant infections caused by VRE and the ESKAPE pathogens.

## RESULTS

### Multiple strains of *S. mutans* reduce viability of Gram-positive pathogens in co-culture

*E. faecalis* is found in human oral cavities (25, 27, 34–36) and is commonly found in root canal infections (37, 38). Therefore, we asked whether *E. faecalis* and oral Streptococci compete with each other. We co-cultured *Streptococcus gordonii* DL1, *Streptococcus sanguinis* SK36, and *S. mutans* UA159 with *E. faecalis* strain OG1RF, a derivative of a human oral isolate (27). We performed the co-cultures using a 1:10 *E. faecalis*:*Streptococcus* spp. inoculum ratio in Brain-Heart Infusion (BHI) broth and quantified CFU/mL of *E. faecalis* OG1RF at t = 0, 2, 4, 6, and 24 h. OG1RF CFU/mL increased over the course of 24 h when grown in BHI monoculture, and co-culture with *S. gordonii* and *S. sanguinis* did not affect OG1RF growth (**Fig. 1A**). However, OG1RF growth was significantly decreased at 4 and 6 h of co-culture with *S. mutans* UA159 compared to the inoculum of 10^7^ CFU/mL at 0 h, indicating that co-culture with UA159 is bactericidal toward OG1RF (**Fig. 1A**). This is consistent with a previous observation that UA159 inhibits *E. faecalis* on solid media (44). At 24 h of co-culture with UA159, we observed an increase in OG1RF CFU/mL to near inoculum levels and a 3.6 log increase compared to the 6 h timepoint (**Fig. 1A**), suggesting that the OG1RF population is able to recover from killing. We then performed co-cultures of OG1RF with *S. mutans* UA159 at inoculation ratios between 1:1 and 1:10 and found that UA159 killing of OG1RF is dose-dependent (**Fig. S1A**). Because we observed the largest effect with a 1:10 ratio of OG1RF:UA159, we proceeded to use this ratio for subsequent co-culture experiments.

**Fig. 1.**
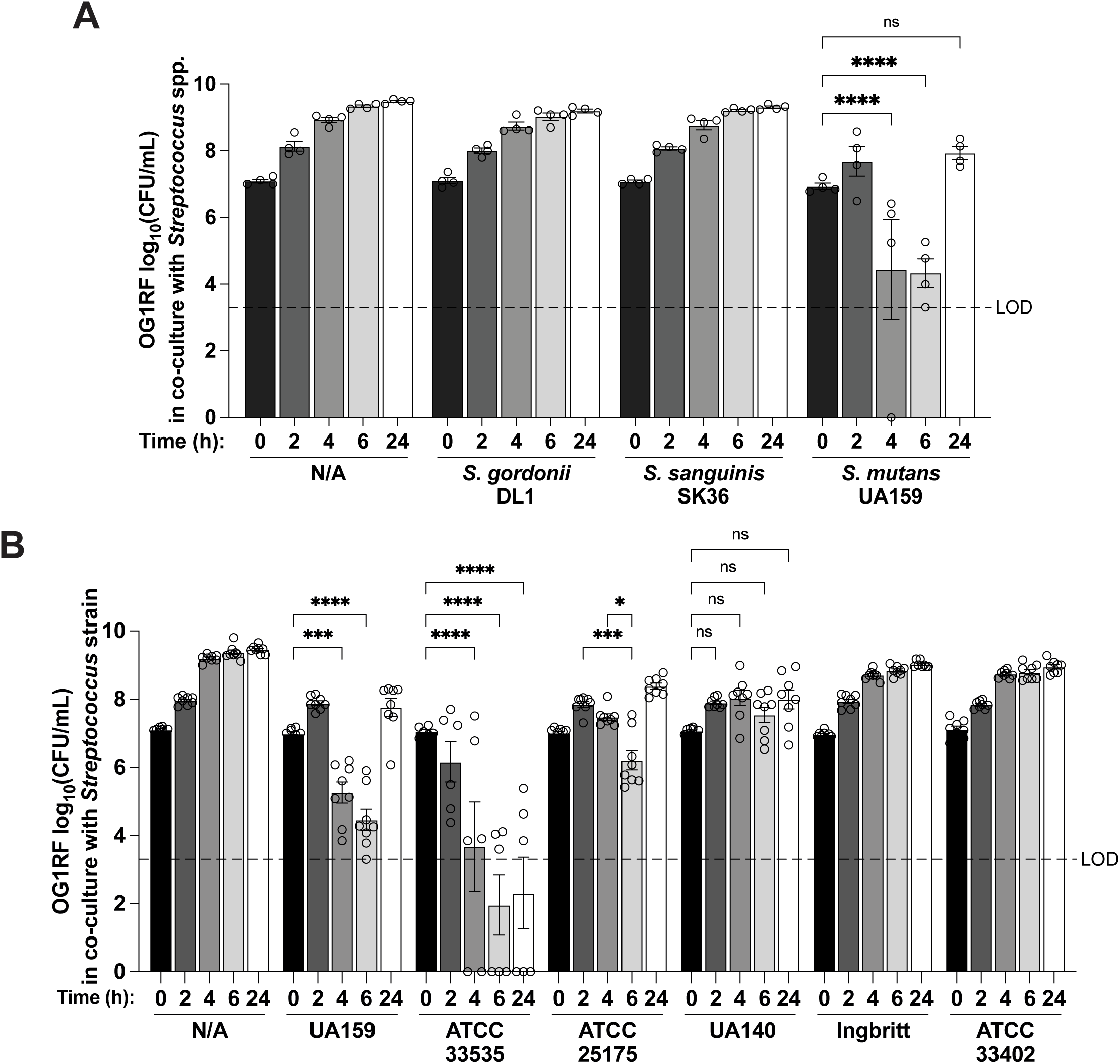
*S. mutans* reduces viability of *E. faecalis* OG1RF in co-culture. **A.** *E. faecalis* OG1RF CFU/mL in 1:10 co-culture with indicated *Streptococcus* spp. strain in BHI at 0, 2, 4, 6, and 24 h. For each culture, n = 2 technical replicates for each of n = 4 biological replicates. **B.** *E. faecalis* OG1RF CFU/mL in 1:10 co-culture with indicated *S. mutans* strain or *S. sobrinus* ATCC 33402 in BHI at 0, 2, 4, 6, and 24 h. For each culture, n = 2 technical replicates for each of n = 6-8 biological replicates. For A and B, two-way ANOVA was used for statistical analysis (ns, not significant; *, p = 0.0162; ***, p ≤ 0.0006; ****, p < 0.0001), each dot represents the mean of technical replicates, bars represent the mean of biological replicates, error bars represent standard errors of the mean, dashed lines indicate limit of detection (LOD), and data points at y = 0 indicate no CFUs were detected.

Next, we investigated whether killing of OG1RF is specific to *S. mutans* UA159 or if other strains of *S. mutans* have inhibitory activity. We performed co-culture experiments with OG1RF and five strains of *S. mutans*: UA159, UA140, ATCC 25174, ATCC 33535, and Ingbritt. We also included *Streptococcus sobrinus* strain ATCC 33402 since it was previously classified as an *S. mutans* strain. Of the six strains tested, UA159, ATCC 25175, and ATCC 33535 significantly reduced CFU/mL of OG1RF (**Fig. 1B**). UA140 did not reduce viable OG1RF cells below that of the inoculum, but OG1RF growth was inhibited and did not significantly increase over time (**Fig. 1B**). We observed differences in the timing and magnitude of killing by UA159, ATCC 25175, and ATCC 33535. While co-culture with UA159 reduced OG1RF CFU/mL starting at 4 h and by a maximum of 3.4 logs at 6 h compared to the highest CFU/mL at 2 h, co-culture with ATCC 25175 only reduced OG1RF CFU/mL by 1.6 logs at 6 h compared to 2 h and OG1RF viability recovered at 24 h (**Fig. 1B**). Co-culture with ATCC 33535, however, reduced OG1RF CFU/mL by 2 h and reduced CFU/mL by a maximum of 5.1 logs at 6 h compared to 0 h (**Fig. 1B**). By 24 h of co-culture with ATCC 33535, OG1RF had not recovered from killing (**Fig. 1B**). Additionally, co-culture with ATCC 33535 eliminated any observable OG1RF CFUs for half of biological replicates at 4, 6, and 24 h (**Fig. 1B**). Therefore, while multiple strains of *S. mutans* are able to kill or inhibit OG1RF, there are interesting differences in the timing and magnitude of killing, suggesting differences in *S. mutans* killing mechanisms and OG1RF resistance to killing.

We then asked if *S. mutans* UA159 is able to kill other strains of *E. faecalis* and performed co-culture experiments with eleven other *E. faecalis* clinical isolates (listed in **Table 1**). To facilitate selection of UA159 for CFU enumeration, we constructed a strain of UA159 with an erythromycin resistance cassette integrated at a neutral chromosomal site (UA159 erm^R^) and used this strain for subsequent co-culture experiments. When co-cultured with UA159 erm^R^, all eleven *E. faecalis* strains tested had lower CFU/mL at 4 h compared to 0 h, including the vancomycin-resistant strain V583 (**Fig. 2A**). We then expanded our panel to include *Enterococcus faecium* and *Enterococcus hirae* isolates. Again, when co-cultured with UA159 erm^R^, viability of all *Enterococcus* isolates was reduced at 4 h compared to 0 h (**Fig. 2B**). None of the *Enterococcus* isolates tested had growth defects in BHI monoculture (**Fig. S1B,C**). We further expanded our panel to include both Gram-positive and Gram-negative pathogens, with a representative from each member of the ESKAPE pathogens. In addition to the *Enterococcus* spp. strains tested, *S. aureus* and *S. epidermidis* were also killed by UA159 erm^R^ by 4 h of co-culture (**Fig. 2C**). The growth of the Gram-negative pathogens (*Klebsiella pneumoniae*, *Acinetobacter baumannii, Pseudomonas aeruginosa,* and *Enterobacter aerogenes*) was not impacted by co-culture with UA159 erm^R^ (**Fig. 2C**). Altogether, these data suggest that *S. mutans* UA159 kills a broad range of Gram-positive pathogens, but this killing does not extend to Gram-negative bacteria.

**Table 1.**
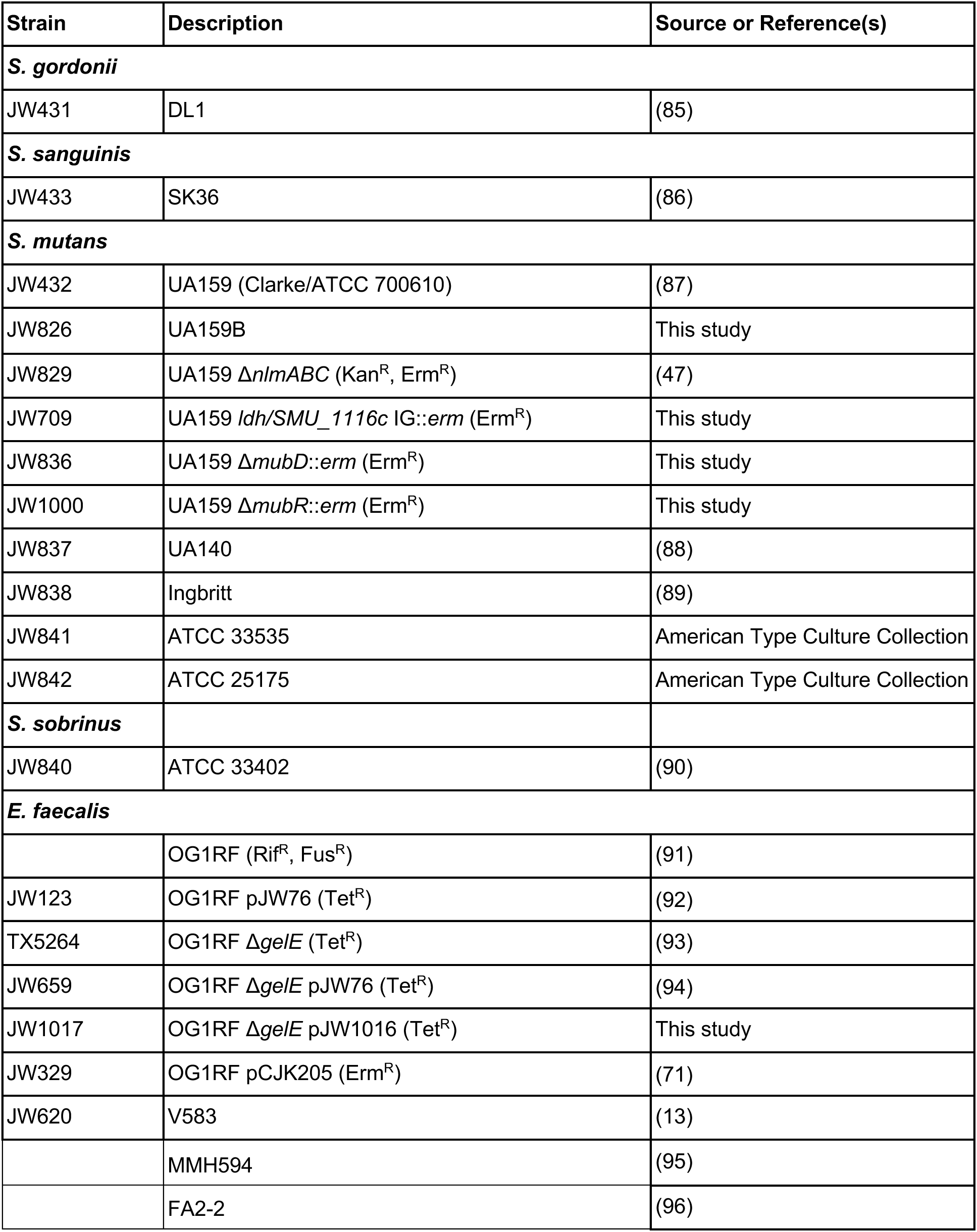
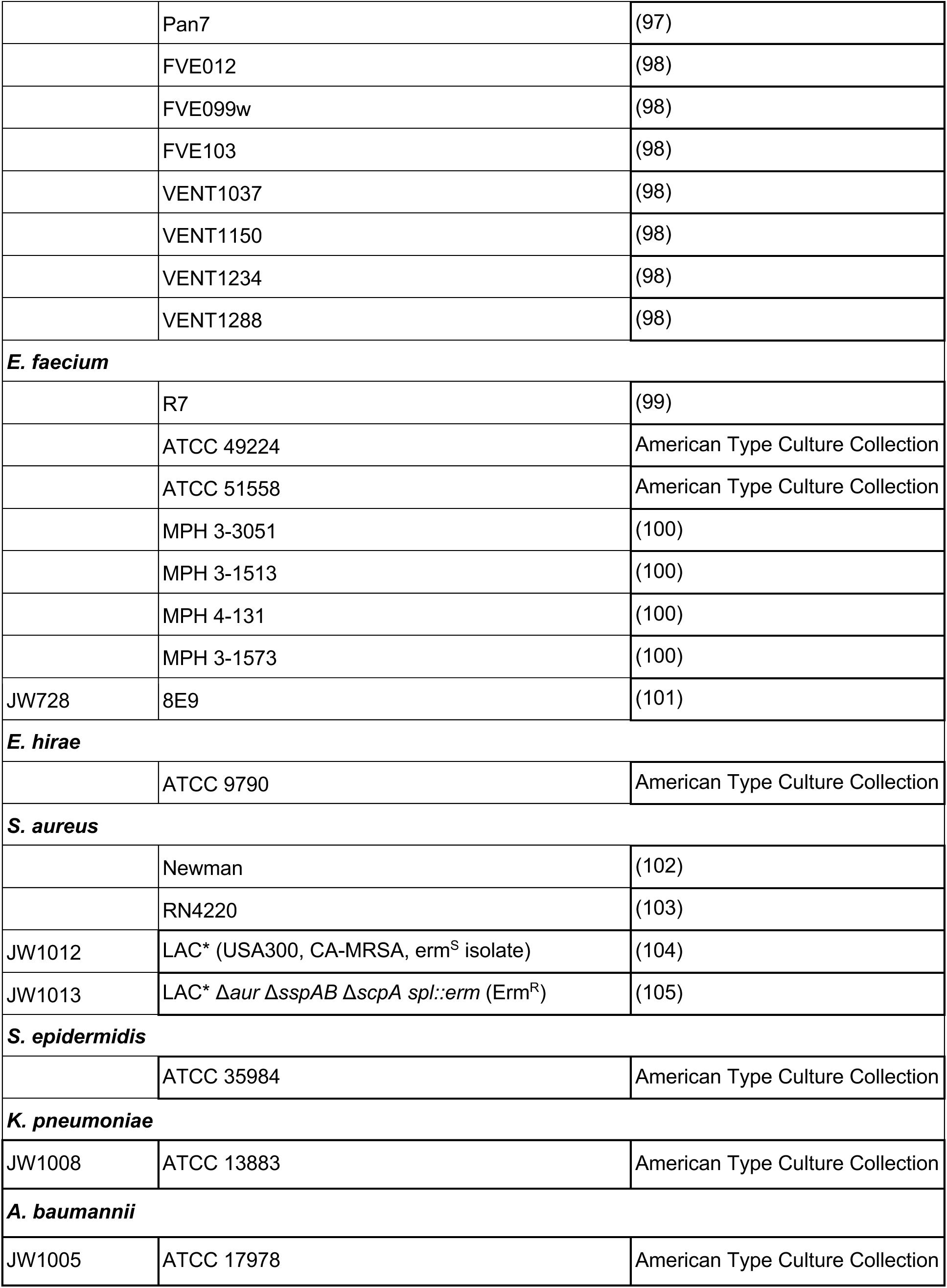
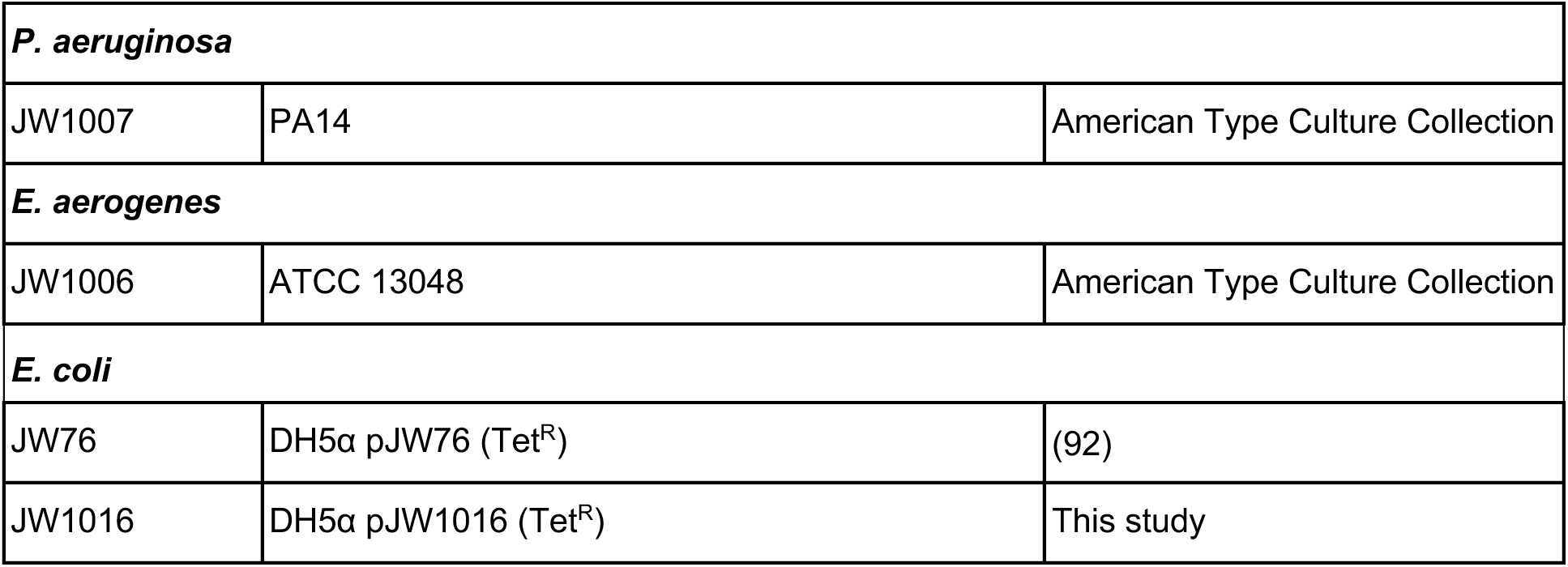
Strains used in this study.

**Fig. 2.**
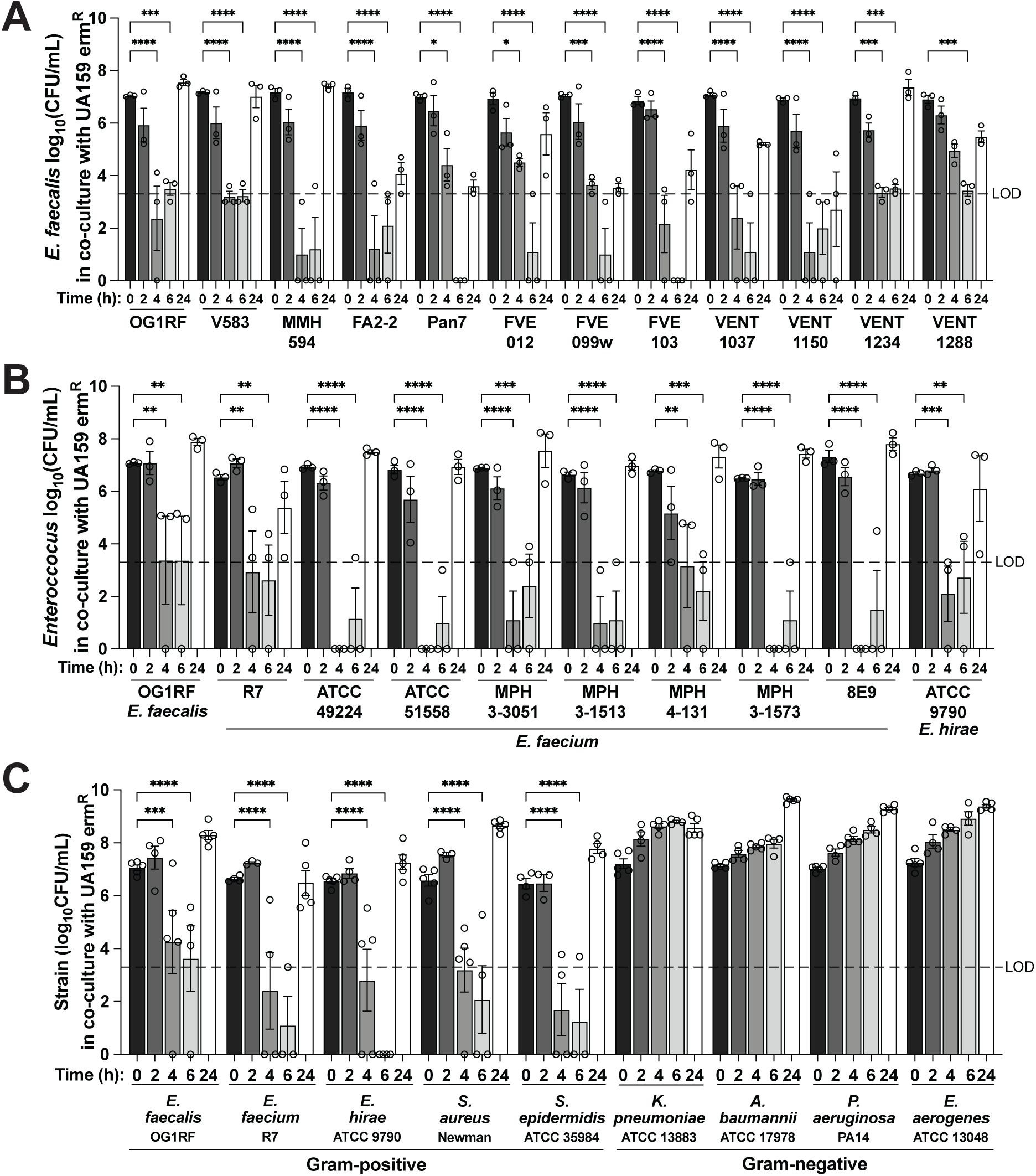
*S. mutans* UA159 kills Gram-positive clinical isolates including ESKAPE pathogens. **A.** CFU/mL of indicated *E. faecalis* clinical isolates in 1:10 co-culture with *S. mutans* UA159 erm^R^ in BHI at 0, 2, 4, 6, and 24 h. For each culture, n = 2 technical replicates for each of n = 3 biological replicates. **B.** CFU/mL of indicated *Enterococcus* spp. strains in 1:10 co-culture with *S. mutans* UA159 erm^R^ in BHI at 0, 2, 4, 6, and 24 h. For each culture, n = 2 technical replicates for each of n = 3 biological replicates. **C.** CFU/mL of indicated strains in 1:10 co-culture with *S. mutans* UA159 erm^R^ in BHI at 0, 2, 4, 6, and 24 h. For each culture, n = 2 technical replicates for each of n = 4-5 biological replicates. For A, B, and C, two-way ANOVA was used for statistical analysis (*, p < 0.03; **, p ≤ 0.009 ***; p ≤ 0.0008; ****, p < 0.0001), each dot represents the mean of technical replicates, bars represent the mean of biological replicates, error bars represent standard errors of the mean, dashed lines indicate limit of detection (LOD), and data points at y = 0 indicate no CFUs were detected.

To begin to probe the mechanism underlying UA159 killing of Gram-positive species, we first tested whether UA159 altered culture pH such that co-culture strains were not able to survive, as *S. mutans* and other lactic acid bacteria (like *Enterococcus* spp.) reduce pH during growth (45). We measured the pH of 24 h monocultures of OG1RF and UA159 as well as an OG1RF/UA159 co-culture. Compared to uninoculated BHI, OG1RF decreased the culture pH to 5.3, UA159 decreased the culture pH to 5.0, and the OG1RF/UA159 co-culture pH matched that of a UA159 monoculture (**Fig. S2A**). We grew OG1RF in BHI at a range of pH values from 2-7 and determined that OG1RF viability was not reduced above pH 4 (**Fig. S2B**). Therefore, it is unlikely that acid production by UA159 is responsible for killing of OG1RF. *S. mutans* UA159 produces two non-lantibiotic mutacins, mutacin IV and mutacin V, encoded by *nlmAB* and *nlmC*, respectively (46). Since UA159 production of mutacin IV and V inhibits the growth of various *Streptococcus* spp. (47), we hypothesized that mutacin production by UA159 is responsible for killing of OG1RF. However, when we co-cultured OG1RF with UA159 lacking genes for both mutacins, OG1RF growth was not different from OG1RF co-cultured with UA159 erm^R^ (**Fig. S2C**), leading to the conclusion that mutacin production by UA159 is not responsible for killing of OG1RF.

### *E. faecalis* gelatinase activity promotes resistance to killing by *S. mutans* UA159

At 24 h of co-culture with *S. mutans* UA159, we observed OG1RF recovery from killing, and 24 h CFU/mL values were similar to the inoculum (**Fig. 1A**). The CFU/mL of UA159 in co-culture with OG1RF did not decrease over 24 h (**Fig. S2D**). Therefore, the increase of OG1RF CFU/mL at 24 h was not due to fewer UA159 cells in co-culture. To test whether OG1RF acquires lasting resistance to killing by UA159, we isolated OG1RF colonies from a 24 h co-culture with UA159 and reinoculated them into a new co-culture with UA159. The 24 h co-culture isolates were killed by UA159 to the same degree as the OG1RF parent strain (**Fig. S2E**), suggesting OG1RF recovery is not the result of any mutations that may have occurred during co-culture and instead may be due to an OG1RF factor that promotes resistance.

While testing the panel of *E. faecalis* isolates shown in **Fig. 2A**, we noticed that several isolates in addition to OG1RF recovered from killing by 24 h of co-culture, but some did not recover by 24 h. One of the *E. faecalis* strains that does not recover from killing is FA2-2 (**Fig. 2A**), which does not have gelatinase activity due to a defect in the *fsr* quorum sensing system that controls the expression of GelE, an extracellular protease that is a key virulence factor for *E. faecalis* (48–51). Conversely, strains known to have gelatinase activity, OG1RF and V583 (49, 52), did recover from killing (**Fig. 2A**). This led us to hypothesize that gelatinase activity of *E. faecalis* predicts recovery from killing by UA159. To determine which of the *E. faecalis* strains we tested in co-culture with UA159 have gelatinase activity, we used a gelatin plate-based assay to detect gelatinase activity after overnight growth. Of the twelve *E. faecalis* strains assayed, five displayed gelatinase activity as indicated by an opaque halo around the site where strains were spotted (**Fig. 3A**). Four of the gelatinase-positive strains, OG1RF, MMH594, VENT1234, and V583, recovered from killing at or above the level of inoculum, while all gelatinase-negative strains did not recover from killing (**Fig. 3A**). FVE012 is gelatinase-positive, and although it did not recover to the level of inoculum, it did recover more than the gelatinase-negative strains (**Fig. 3A**). We quantified gelatinase activity by measuring the radius of the opaque halo surrounding the strains spotted on gelatin plates and found that FVE012 had reduced gelatinase activity compared to the other gelatinase-positive strains (**Fig. S3A**). Therefore, decreased gelatinase expression may explain the intermediate recovery of FVE012 from killing. Indeed, when we plotted average 24 h co-culture CFU/mL and average gelatinase halo radius for each of the *E. faecalis* strains, strains with any gelatinase activity recovered from killing more than strains without gelatinase activity (**Fig. S3A**). Together, our gelatinase assay and co-culture results suggest that *E. faecalis* gelatinase activity predicts recovery from killing by UA159. This observation is also consistent with the timing of GelE activity reaching its peak when growth reaches stationary phase (53).

**Fig. 3.**
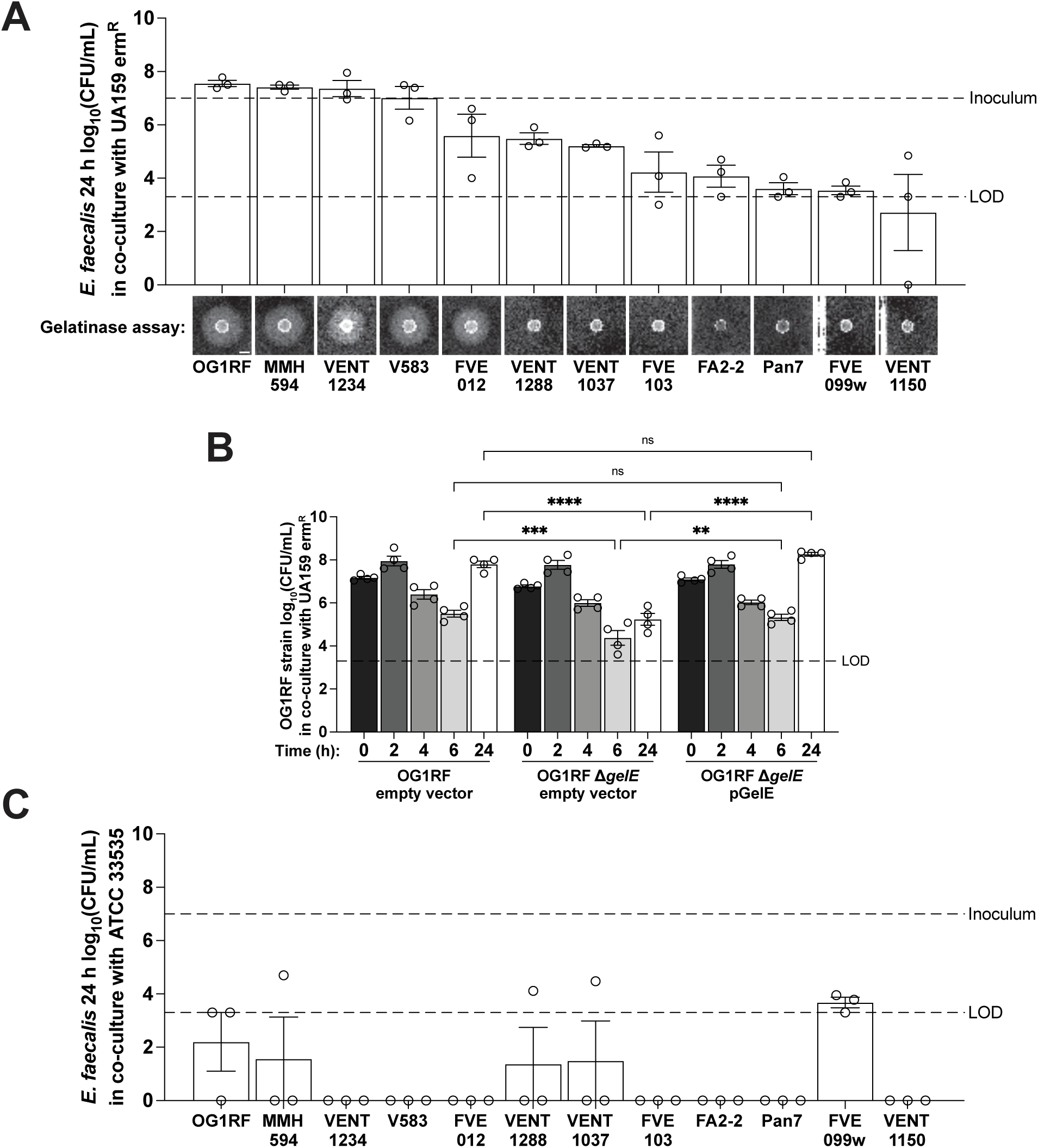
*E. faecalis* gelatinase activity is required for resistance to killing by *S. mutans* UA159. **A.** CFU/mL of indicated *E. faecalis* clinical isolates in 1:10 co-culture with *S. mutans* UA159 erm^R^ in BHI at 24 h (data is the same as shown in Fig. 2A). Representative gelatinase assay images are included on the x-axis for each *E. faecalis* clinical isolate. For each co-culture, n = 2 technical replicates for each of n = 3 biological replicates. For the gelatinase assays, n = 2 technical replicates per n = 2-5 biological replicates. Scale bar represents 5 mm. **B.** CFU/mL of indicated OG1RF strains in 1:10 co-culture with *S. mutans* UA159 erm^R^ in BHI at 0, 2, 4, 6, and 24 h. For each culture, n = 2 technical replicates for each of n = 4 biological replicates. Two-way ANOVA was used for statistical analysis (**, p = 0.0015; ***, p = 0.0002; ****, p < 0.0001). **C.** CFU/mL of indicated *E. faecalis* clinical isolates in 1:10 co-culture with *S. mutans* ATCC 33535 in BHI at 24 h. For each co-culture, n = 2 technical replicates for each of n = 3 biological replicates. For A and C, upper dashed lines indicate the *E. faecalis* inoculum of 10^7^ CFU/mL. For A, B, and C, each dot represents the mean of technical replicates, bars represent the mean of biological replicates, error bars represent standard errors of the mean, lower dashed lines indicate limit of detection (LOD), and data points at y = 0 indicate no CFUs were detected.

To determine the requirement of gelatinase for recovery from killing by UA159, we co-cultured a strain of OG1RF lacking the GelE gelatinase with UA159. While the OG1RF Δ*gelE* strain did not have a growth defect in monoculture (**Fig. S3**), OG1RF Δ*gelE* had significantly fewer CFU/mL at 24 h of co-culture than the OG1RF parental strain, and expression of *gelE* from an inducible plasmid restored the OG1RF Δ*gelE* recovery defect (**Fig. 3B**). These results demonstrate that gelatinase activity is required for full *E. faecalis* recovery from killing by UA159. Interestingly, the 12 *E. faecalis* isolates do not recover from killing by ATCC 33535 by 24 h (**Fig. 3C**), suggesting that the mechanism of inhibition by this strain of *S. mutans* differs from that of UA159.

We next asked whether extracellular proteases could protect other Gram-positive bacteria from killing by UA159, given that the *S. aureus* strains Newman, RN4220, and LAC* recovered from killing by 24 h (**Fig. 2C** and **Fig. S3CD**). *S. aureus* produces several extracellular proteases (54, 55), so we wondered if any of these enzymes contribute to recovery from killing by UA159. We performed co-cultures with UA159 and the *S. aureus* MRSA strain LAC* lacking all four known extracellular proteases and found that the LAC* protease knockout strain recovered from killing similarly to the parental LAC* strain (**Fig. S3D**). This result demonstrates that *S. aureus* recovery from killing by UA159 is not mediated by the production of these extracellular proteases and that mechanisms of killing and/or recovery from killing likely vary depending on the target species.

### *S. mutans* UA159 mutanobactin biosynthesis genes are required to kill Gram-positive pathogens

During the course of our co-culture experiments, we tested a different stock of UA159 that was obtained from another lab. Interestingly, this stock of UA159 did not kill OG1RF in co-culture, and growth of OG1RF in co-culture with this strain was similar to that of OG1RF in monoculture (**Fig. 4A**). Because of this difference in killing phenotype, we refer to this strain as UA159B. To determine why UA159B was unable to reduce the number of viable OG1RF cells in co-culture, we sequenced the genome of UA159B and compared it to the original isolate of UA159. This revealed an insertion mutation in UA159B *mubD* by ISSmu1, an IS3 family transposable element (**Fig. 4B**). The *mubD* gene is a part of the biosynthetic gene cluster that produces mutanobactin, a hybrid polyketide/nonribosomal peptide (56, 57) that promotes oxygen and hydrogen peroxide stress tolerance (57–59), represses the yeast-mycelium morphological transition of the oral fungal pathogen *Candida albicans* (56, 60), and increases immunomodulatory activity (58). Mutanobactin has not, however, previously been shown to have antibacterial activity against Gram-positive bacteria.

**Fig. 4.**
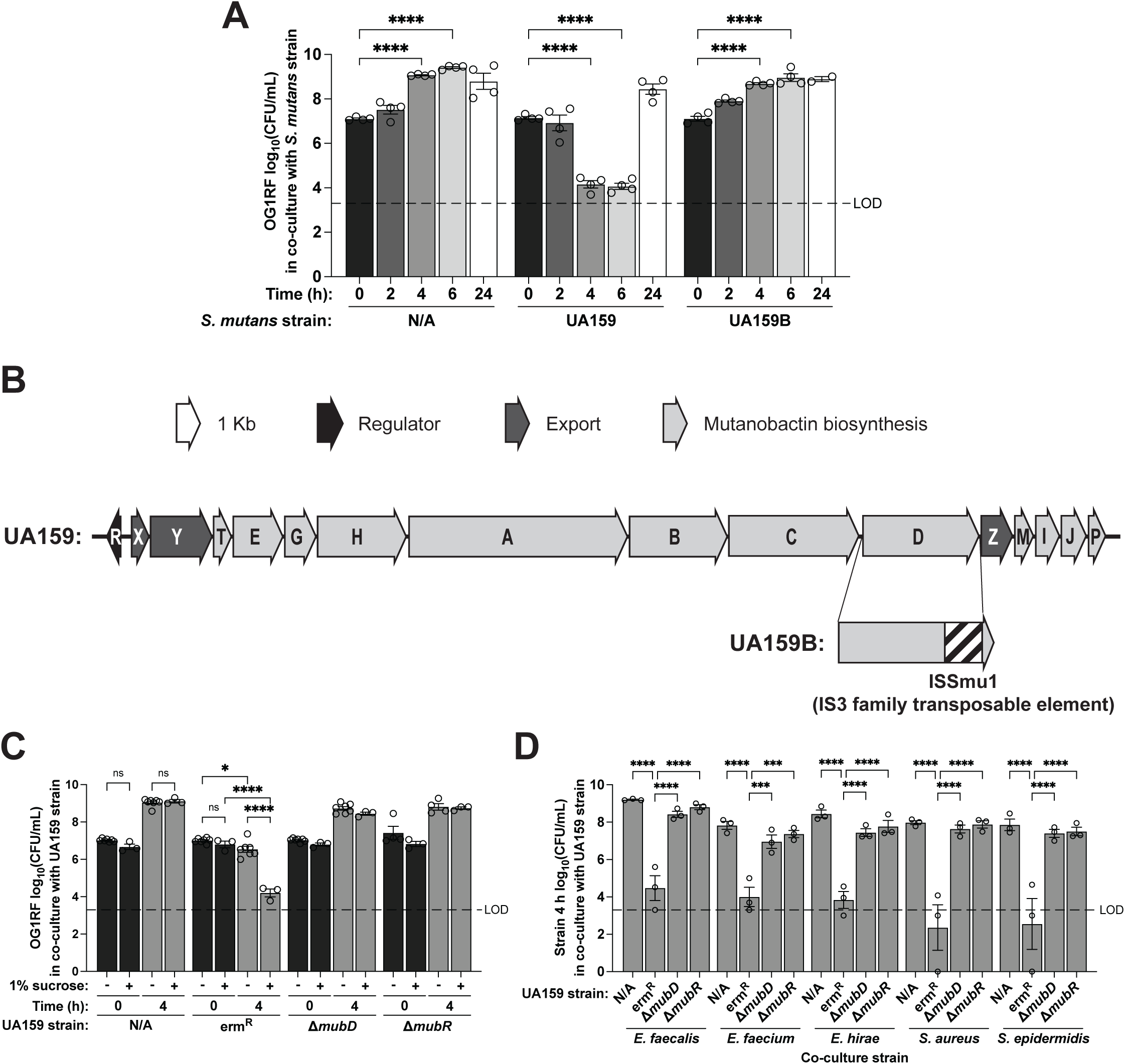
*S. mutans* UA159 mutanobactin genes are required to kill Gram-positive pathogens in co-culture. **A.** *E. faecalis* OG1RF CFU/mL in 1:10 co-culture with indicated *S. mutans* strain in BHI at 0, 2, 4, 6, and 24 h. For each culture, n = 2 technical replicates for each of n = 2-4 biological replicates. **B.** Genetic organization of the *S. mutans* UA159 mutanobactin biosynthetic gene cluster and location of the UA159B ISSmu1 insertion in *mubD*. **C.** *E. faecalis* OG1RF CFU/mL in 1:10 co-culture with indicated *S. mutans* UA159 strain in BHI ± 1% sucrose at 0 and 4 h. For each culture, n = 2 technical replicates for each of n = 3-7 biological replicates. **D.** CFU/mL of indicated strains in 1:10 co-culture with indicated *S. mutans* UA159 strain in BHI at 4 h. For each culture, n = 2 technical replicates for each of n = 3 biological replicates. For A, C, and D, two-way ANOVA was used for statistical analysis (ns, not significant; *, p = 0.0153; **, p ≤ 0.009; ***, p ≤ 0.0008; ****, p < 0.0001), each dot represents the mean of technical replicates, bars represent the mean of biological replicates, error bars represent standard errors of the mean, dashed lines indicate limit of detection (LOD), and data points at y = 0 indicate no CFUs were detected.

To confirm that the mutation in *mubD* is responsible for the inability of UA159B to kill OG1RF, we deleted *mubD* in our original UA159 strain background and co-cultured the Δ*mubD* strain with OG1RF. UA159 Δ*mubD* did not kill OG1RF in co-culture (**Fig. 4C, Fig. S4A**), confirming that *mubD* is required for UA159 antibacterial activity. We also generated a deletion of *mubR*, which encodes a TetR family transcription factor that activates expression of the *mub* gene cluster. This UA159 Δ*mubR* mutant was also unable to kill OG1RF in co-culture (**Fig. 4C, Fig. S4A**). We examined the genomes of the *S. mutans* strains shown in **Fig. 1B** and identified a putative mutanobactin biosynthetic gene cluster in ATCC 25175. Strain UA140 also encodes for mutanobactin production (57). In our co-cultures, UA140 prevented growth of OG1RF and OG1RF viability was reduced by ATCC 25175 with recovery from killing by 24 h (**Fig. 1B**). This further supports the hypothesis that mutanobactin mediates interbacterial competition between *S. mutans* and other Gram-positive bacteria.

We next wanted to determine whether mutanobactin mediates interbacterial competition in conditions more relevant to the oral cavity. Sucrose promotes the formation of dental caries by lowering pH and serving as a substrate for extracellular and intracellular polysaccharide synthesis (45, 61–66). To see if mutanobactin-mediated killing by UA159 occurred in the presence of this physiologically relevant sugar, we performed co-culture assays between UA159 and OG1RF in BHI supplemented with 1% sucrose. At 4 h of co-culture with UA159, we observed enhanced killing of OG1RF in BHI + 1% sucrose compared to unsupplemented BHI (**Fig. 4C**). Co-culture with either the UA159 Δ*mubD* or Δ*mubR* mutant in BHI + 1% sucrose did not reduce OG1RF CFU/mL compared to OG1RF monoculture (**Fig. 4C, Fig. S4B**), indicating mutanobactin is required for killing in the presence of sucrose. Finally, we co-cultured the UA159 Δ*mubD* and Δ*mubR* mutants with representative Gram-positive pathogens. Both *mub* genes were required for UA159 to kill each of the Gram-positive pathogens tested (**Fig. 4D, Fig. S4C-D**). Taken together, these results provide strong genetic evidence that *S. mutans* antibacterial activity toward Gram-positive species is dependent on the production of mutanobactin, and that mutanobactin-mediated competition could occur in sucrose-rich conditions in the oral cavity.

### Mutanobactin-mediated killing increases target cell membrane permeability and is contact-dependent

Mutanobactin is a cyclic lipopeptide that is similar in structure to daptomycin, a lipopeptide antibiotic that targets Gram-positive bacterial cell membranes (56, 57, 60, 67). Daptomycin exerts its antibacterial effect by inserting into the Gram-positive bacterial cell membrane, resulting in membrane depolarization (68, 69) Since mutanobactin and daptomycin share structural similarities, we hypothesized that mutanobactin also decreases Gram-positive cell viability by targeting the bacterial cell membrane. To test this, we performed cell membrane permeability assays using chlorophenyl red-β-D-galactopyranoside (CPRG) to measure OG1RF cell membrane integrity (70, 71). In this assay, β-galactosidase activity of LacZ constitutively expressed in OG1RF is assessed by measuring the absorbance of chlorophenyl red, the cleavage product of the cell membrane-impermeable compound CPRG. We cultured OG1RF constitutively expressing *lacZ* from the pCJK205 plasmid in BHI with subinhibitory concentrations of daptomycin and ciprofloxacin, a fluoroquinolone antibiotic that functions by inhibiting DNA replication (72). As expected, OG1RF cell membrane permeability relative to the untreated control was significantly increased when grown with daptomycin, but not with ciprofloxacin (**Fig. 5A**). When we co-cultured OG1RF pCJK205 with UA159, OG1RF cell membrane permeability was significantly increased compared to growth in monoculture, and this increase was dependent on UA159 *mubD* and *mubR* (**Fig. 5B**). These results support our hypothesis that UA159 production of mutanobactin reduces target cell viability by disrupting cell membrane integrity.

**Fig. 5.**
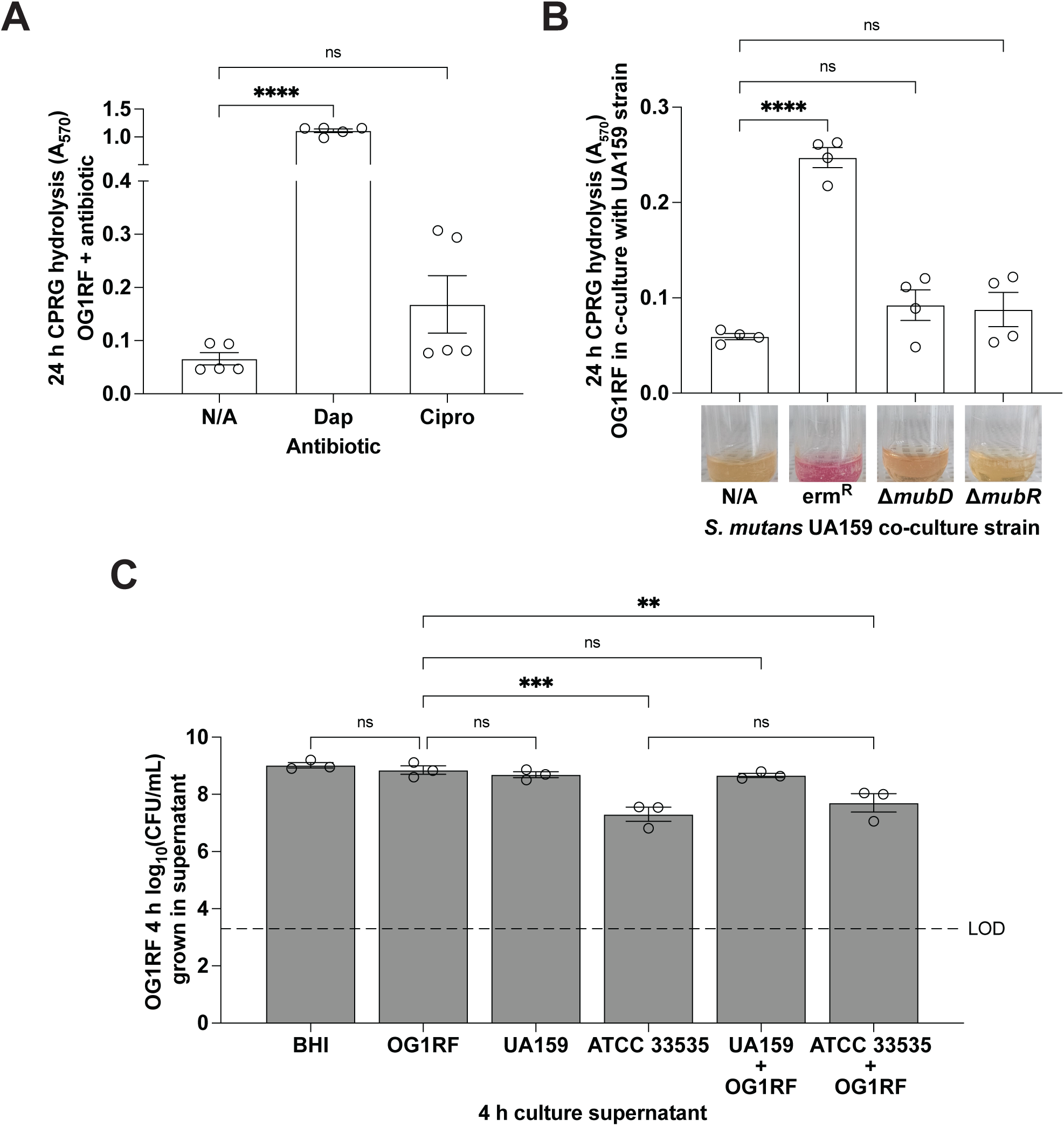
Mutanobactin production by *S. mutans* UA159 increases cell membrane permeability in a contact-dependent manner. **A.** CPRG hydrolysis of *E. faecalis* OG1RF at 24 h culture in TBS-D containing subinhibitory concentrations of indicated antibiotic. For each culture, n = 2 technical replicates for each of n = 5 biological replicates. **B.** CPRG hydrolysis of *E. faecalis* OG1RF at 24 h co-culture with indicated *S. mutans* UA159 strain in TBS-D. Representative images of CPRG assay cultures are included on the x-axis for each co-culture. For each culture, n = 2 technical replicates for each of n = 4 biological replicates. **C.** *E. faecalis* OG1RF CFU/mL grown in BHI containing 50% cell-free supernatant from indicated 4 h monoculture or co-culture. For each culture, n = 2 technical replicates for each of n = 3 biological replicates, dashed line indicates limit of detection (LOD), and data points at y = 0 indicate no CFUs were detected. For A-C, one-way ANOVA was used for statistical analysis (ns, not significant; **, p = 0.0053; ** p = 0.0096; ***, p = 0.0009; ****, p < 0.0001), each dot represents the mean of technical replicates, bars represent the mean of biological replicates, error bars represent standard errors of the mean.

We next tested whether mutanobactin-mediated killing by UA159 is contact-dependent by growing OG1RF in UA159 cell-free supernatant. There was no difference in OG1RF growth when grown in supernatants derived from OG1RF or UA159 monocultures (**Fig. 5C**). We next considered the possibility that UA159 only produces mutanobactin in the presence of other microbes but did not observe inhibition of OG1RF when OG1RF was grown in cell-free supernatant from a 4 h 1:10 co-culture of OG1RF and UA159 (**Fig. 5C**). Thus, mutanobactin-mediated antibacterial activity by UA159 is likely contact-dependent. Interestingly, when we grew OG1RF in cell-free supernatant from the non-mutanobactin-producing strain ATCC 33535, we observed a 1.7 log inhibition of OG1RF CFU/mL and similar inhibition when OG1RF was grown in cell-free supernatant from OG1RF/ATCC 33535 co-cultures (**Fig. 5C**). While OG1RF viability in ATCC 33535 cell-free supernatant was not reduced to the same extent as when co-cultured with ATCC 33535 cells (**Fig. 1B**), our data suggest that ATCC 33535 inhibition of OG1RF does not require cell-cell contact. This is in contrast to the contact-dependent mechanism of UA159 mutanobactin-mediated killing. In support of this hypothesis, *E. faecalis* isolates do not recover by 24 h of co-culture with ATCC 33535, regardless of whether they produce gelatinase (**Fig. 3C**). This suggests ATCC 33535 kills *E. faecalis* using a mechanism and natural product different from that of UA159.

Mutanobactin production by *S. mutans* UA159 negatively impacts *E. faecalis* biofilms.

*S. mutans* is typically found within dental plaque biofilms on the surface of teeth (45). Therefore, we sought to investigate interactions between UA159 and OG1RF in the context of biofilms. We first tested whether OG1RF is able to form biofilms when co-cultured with UA159. For these assays, we co-cultured OG1RF and UA159 in the presence of an Aclar fluoropolymer disk and quantified CFU/mL in the planktonic phase as well as the number of biofilm CFUs per Aclar disk at 4 and 24 h (**Fig. 6A**). When inoculated in a 1:1 ratio with UA159, OG1RF biofilm viability decreased at 24 h, and OG1RF growth decreased at both 4 and 24 h when inoculated 1:10 with UA159 (**Fig. 6B**). Planktonic OG1RF CFU/mL were also reduced when co-cultured with UA159 (**Fig. S5A**). Next, we determined whether mutanobactin production by UA159 is required for UA159 inhibition of biofilm formation by performing 1:10 co-cultures of OG1RF with UA159 Δ*mubD* and Δ*mubR* in the presence of an Aclar disk. Neither *mubD* nor *mubR* deletion strains reduced viability of OG1RF biofilms or planktonic cells (**Fig. 6C, Fig. S5B**). Together, these results indicate that UA159 mutanobactin production prevents OG1RF from forming biofilms.

**Fig. 6.**
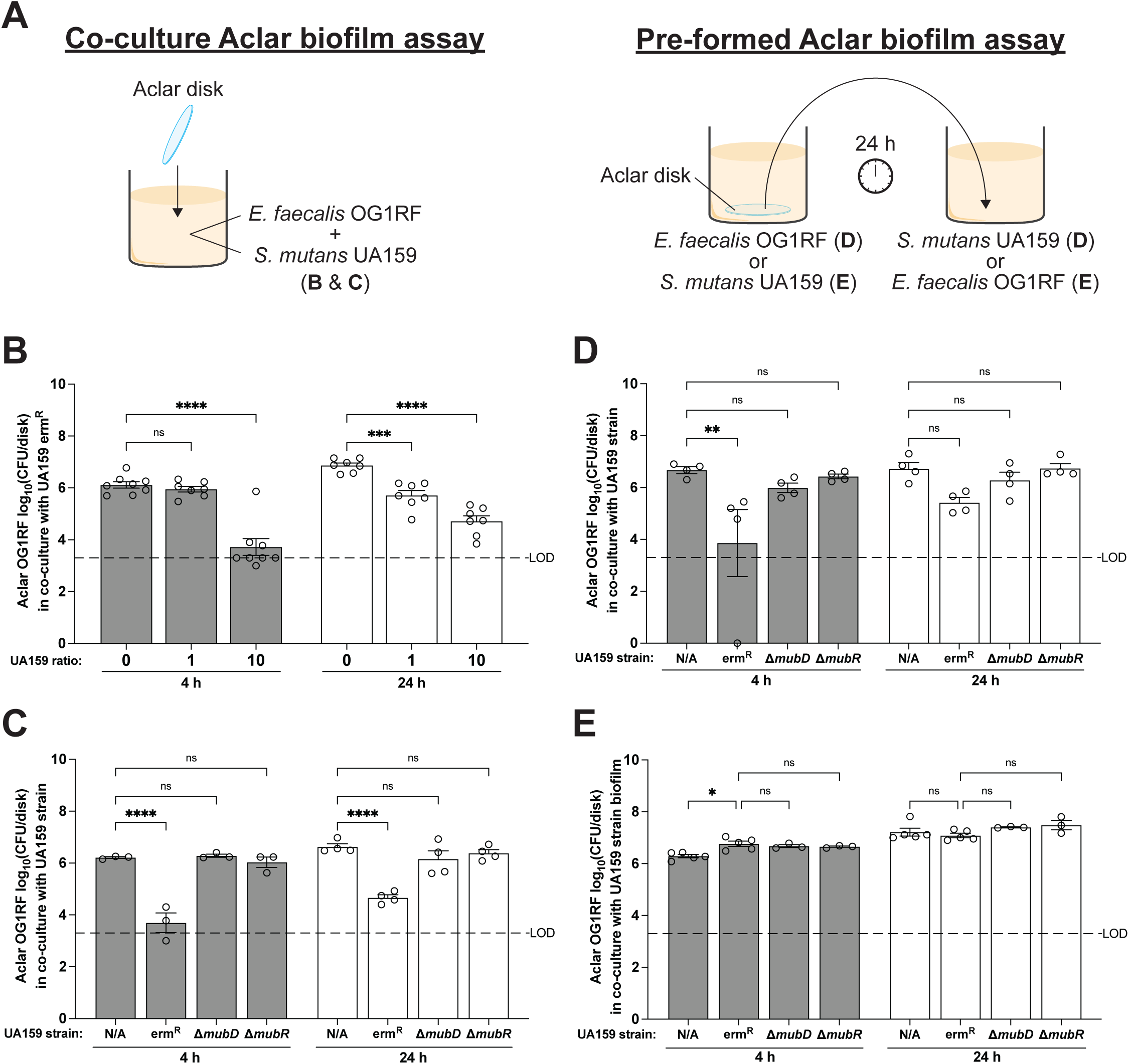
Mutanobactin production by *S. mutans* UA159 prevents *E. faecalis* OG1RF biofilm formation and kills biofilm-contained OG1RF. **A.** Experimental set-ups for the co-culture Aclar biofilm assays for panels B and C and the pre-formed Aclar biofilm assays for panels D and E. For the co-culture biofilm assays, Aclar disks are submerged in media inoculated with both *E. faecalis* OG1RF and *S. mutans* UA159. For the pre-formed biofilm assays, Aclar disks are cultured with either OG1RF or UA159 for 24 h before being transferred to an OG1RF or UA159 mono-culture. **B.** *E. faecalis* OG1RF CFU per Aclar disk in co-culture with *S. mutans* UA159 erm^R^ at indicated inoculum ratio of UA159 erm^R^ to 1 part OG1RF in BHI at 4 and 24 h. For each co-culture, n = 2 technical replicates for each of n = 7-8 biological replicates. **C.** *E. faecalis* OG1RF CFU per Aclar disk in 1:10 co-culture with indicated *S. mutans* UA159 strain in BHI at 4 and 24 h. For each co-culture, n = 2 technical replicates for each of n = 3-4 biological replicates. **D.** *E. faecalis* OG1RF CFU per Aclar disk of OG1RF pre-formed Aclar biofilm in co-culture with indicated *S. mutans* UA159 strain in BHI at 4 and 24 h. For each co-culture, n = 2 technical replicates for each of n = 4 biological replicates. **E.** *E. faecalis* OG1RF CFU per Aclar disk in co-culture with pre-formed Aclar biofilm of indicated *S. mutans* UA159 strain in BHI + 1% sucrose at 4 and 24 h. For each co-culture, n = 2 technical replicates for each of n = 3-5 biological replicates. For B-D, two-way ANOVA was used for statistical analysis (ns, not significant; *, p = 0.0115; **, p = 0.0014; ***, p = 0.0005; ****, p < 0.0001), each dot represents the mean of technical replicates, bars represent the mean of biological replicates, error bars represent standard errors of the mean, dashed lines indicate limit of detection (LOD), and data points at y = 0 indicate no CFUs were detected.

Next, we asked whether UA159 could kill *E. faecalis* cells in pre-formed biofilms. To test this, we grew OG1RF alone in the presence of an Aclar disk for 24 h, transferred the Aclar disk to a culture of UA159, and quantified the number of OG1RF biofilm CFUs per Aclar disk at 4 and 24 h as well as CFU/mL of OG1RF in the planktonic phase (**Fig. 6A**). At 4 h after the transfer of the pre-formed OG1RF Aclar biofilm to a UA159 culture, we observed fewer OG1RF CFUs per Aclar disk and OG1RF planktonic CFU/mL than with OG1RF biofilms transferred to BHI, and the decrease in OG1RF biofilm and planktonic CFUs was dependent on *mubD* and *mubR* (**Fig. 6D, Fig. S5C**). Thus, UA159 production of mutanobactin is able to kill *E. faecalis* biofilms.

Finally, we explored whether pre-formed UA159 biofilms prevent OG1RF biofilm formation by growing UA159 in the presence of an Aclar disk for 24 h, transferring the Aclar disk to a culture of OG1RF, and quantifying the number of OG1RF biofilm CFUs per Aclar disk at 4 and 24 h as well as CFU/mL of OG1RF in the planktonic phase (**Fig. 6A**). While pre-formed UA159 biofilms slightly reduced the number of OG1RF CFU/mL in the planktonic phase by 24 h following UA159 Aclar biofilm transfer (**Fig. S5D**), OG1RF CFUs per Aclar disk were not negatively impacted by the presence of UA159 on the disks, regardless of mutanobactin production (**Fig. 6E**). In sum, these data show that mutanobactin-mediated killing by planktonic UA159 is effective in preventing OG1RF biofilm formation and in inhibiting biofilm-contained OG1RF, but pre-formed UA159 biofilms do not exclude OG1RF from forming dual-species biofilms.

## DISCUSSION

*E. faecalis is* a well-characterized pathobiont that is understudied in the context of the oral cavity despite its prevalence in root canal infections (24–33). Our study aimed to explore microbial factors that could influence *E. faecalis* competition and persistence within the oral cavity. We began by performing co-culture experiments with oral Streptococci, and these co-culture experiments revealed that multiple strains of the oral pathogen *S. mutans* reduce the viability of *E. faecalis* OG1RF (**Fig. 1**). Subsequent experiments demonstrated that *S. mutans* kill other Gram-positive bacteria, including *S. epidermidis* and the ESKAPE pathogen *S. aureus* (**Fig. 2**). We also determined that UA159 inhibitory activity is dependent on the production of mutanobactin, a cyclic lipopeptide that promotes *S. mutans* oxygen and hydrogen peroxide stress tolerance (57–59), inhibits the *C. albicans* yeast-to-mycelium transition (56, 60), and modulates the immune response of human cells (58) (**Fig. 4**). Mutanobactin production increases target cell membrane permeability, similar to the antibiotic daptomycin; and killing by UA159 is contact-dependent (**Fig. 5**). Additionally, we found that an *E. faecalis* virulence factor, GelE, is required for full recovery from killing by UA159, but *E. faecalis* does not recover from killing by ATCC 33535 regardless of GelE (**Fig. 3**). Finally, we assessed UA159 antibacterial activity in the context of biofilms and revealed that UA159 mutanobactin production inhibits OG1RF biofilm formation and kills biofilm-contained OG1RF, while UA159 biofilms do not exclude OG1RF from forming dual-species biofilms (**Fig. 6**). A limitation of our study is that while *E. faecalis* is frequently isolated from the oral cavity, it is not commonly found within dental plaques where *S. mutans* is found (73, 74). Secreted toxins such as the one produced by *S. mutans* ATCC 33535 may diffuse to other niches in the oral cavity, thereby impacting viability and biofilm formation by other bacteria such as *E. faecalis*. Conversely, compounds that are not secreted, such as mutanobactin, may affect *E. faecalis* viability through broader alterations in polymicrobial community structure. The biological significance of mutanobactin within the oral cavity may also be relevant for other Gram-positive bacteria that reside in dental plaque or other oral cavity sites. Future work will determine the spatial and temporal dynamics of *S. mutans* compound production with regard to antibacterial activity toward *E. faecalis* and other oral bacteria. Overall, this work describes a novel function for a previously characterized specialized metabolite, mutanobactin, revealing a new interaction between clinically relevant bacteria that occupy the same ecological niche. We have summarized our results in the context of previous work in the model presented in **Figure 7**. We propose a model where *S. mutans* production of mutanobactin kills *E. faecalis* while *E. faecalis* GelE secretion promotes recovery from killing. *E. faecalis* inhibits *C. albicans* hyphal morphogenesis (75–77), as does mutanobactin (56, 60). *S. mutans* in dental plaque biofilms do not kill *E. faecalis*, which allows *E. faecalis* to persist in the oral cavity and colonize root canals.

**Fig. 7.**
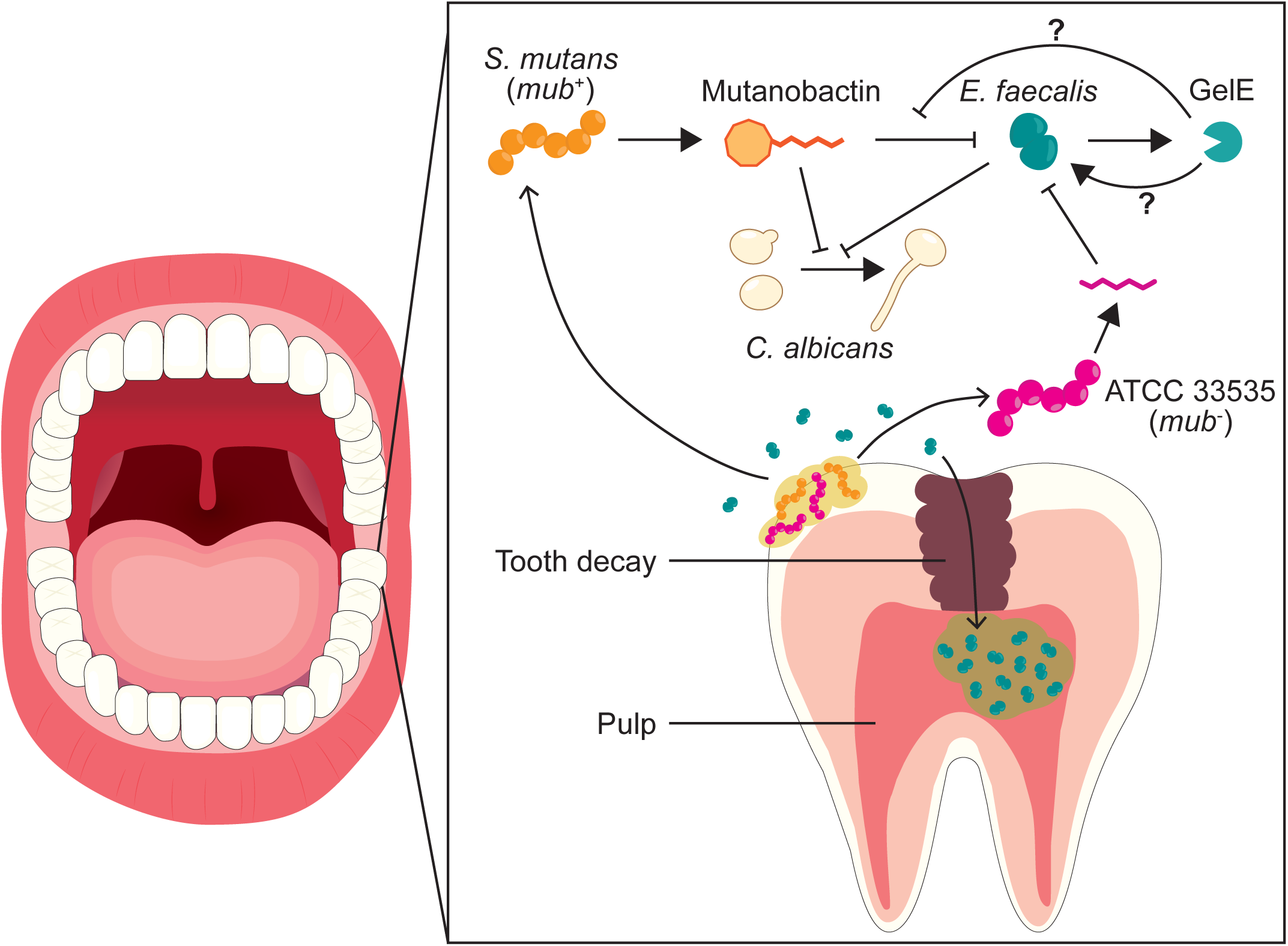
Proposed model of *S. mutans* and *E. faecalis* interactions within the oral cavity. *S. mutans* and *E. faecalis* are both found in the oral cavity and have both been detected in infected root canals. *S. mutans mub*^+^ strains produce mutanobactin, which kill *E. faecalis* in a contact-dependent manner. *E. faecalis* secretion of GelE promotes recovery from killing or prevents mutanobactin from killing. *S. mutans mub*^-^ strain ATCC 33535 does not produce mutanobactin but secretes an uncharacterized compound that kills *E. faecalis* in a contact-independent manner. *C. albicans* hyphal morphogenesis is inhibited by *E. faecalis* (75–77) and by mutanobactin (56, 60). Biofilm-contained *S. mutans* does not inhibit *E. faecalis*, which may allow *E. faecalis* to persist in the presence of *S. mutans* in the oral cavity.

In addition to UA159, *S. mutans* strains UA140 and ATCC 25175 both encode genes for the production of mutanobactin (57). Previous work has shown that the timing and magnitude of mutanobactin gene expression differs between UA159 and UA140, with expression of the UA140 *mub* gene cluster beginning to increase at 7 h of growth and peaking at 10 h opposed to the 4 h beginning and 6 h peak expression observed for UA159 (57). The later timing of UA140 mutanobactin gene expression may explain why we did not observe significant inhibition of OG1RF by UA140 (**Fig. 1C**), as OG1RF *gelE* expression is induced during later stages of growth (53), prior to induction of mutanobactin gene expression. Analysis of the ATCC 25175 genome revealed that its *mub* gene cluster is similar to that of UA159, but differences in *mub* expression levels and/or timing, or other factors, may account for the differences in killing observed between ATCC 25175 and UA159. Whole genome sequencing of ATCC 33535, however, revealed that this strain does not encode genes for the production of mutanobactin, and therefore the mechanism of killing by ATCC 33535 is mutanobactin-independent. This finding highlights the diversity of streptococcal natural products that can mediate interbacterial competition, with contact-dependent and contact-independent mechanisms potentially contributing to differential inhibition of specific microbial targets in specific oral niches.

Without production of the extracellular protease GelE, *E. faecalis* does not fully recover from mutanobactin-mediated killing by 24 h of co-culture with UA159 (**Fig. 3**). While the number of viable GelE-negative cells at 24 h of co-culture with UA159 was 2.1-4.8 logs lower than that of GelE-positive OG1RF, the GelE-negative strains recovered to near the limit of detection by 24 h of co-culture (**Fig. 2A, 3A**). This is consistent with the timing of mutanobactin gene expression in UA159, where expression is at its maximum at 6 h of growth at the end of exponential phase followed by a dramatic decrease in expression once cells enter stationary phase (57). Therefore, *E. faecalis* overcoming antibacterial activity by UA159 is likely influenced by the temporal expression of both enterococcal *gelE* and UA159 mutanobactin. Since expression of *gelE* is controlled by the *fsr* quorum sensing system (53), *gelE* expression should not be activated at early time points in our assays since *E. faecalis* cell density is low when co-cultured with UA159. One possible explanation is that co-culture with UA159 induces *gelE* expression. Another explanation for the requirement of GelE for full recovery is that once mutanobactin production decreases after UA159 reaches stationary phase, GelE facilitates the outgrowth of surviving *E. faecalis* cells. This hypothesis is supported by our data demonstrating that *fsr*-independent induction of *gelE* expression at the beginning of co-culture did not prevent killing of OG1RF at 4 and 6 h of co-culture with UA159 (**Fig. 3C**). Regardless, our GelE results support the hypothesis that *fsr*-induced GelE expression alone is not responsible for *E. faecalis* recovery from killing and it is likely that other factors, streptococcal and/or enterococcal, contribute to *E. faecalis* recovery.

We also observed that other Gram-positive species recover from killing by 24 h of co-culture with UA159 (**Fig. 2C, Fig. S3C**), and that the extracellular proteases of *S. aureus* do not contribute to this recovery (**Fig. S3D**). It is possible that other *S. aureus* factors contribute to recovery from mutanobactin-mediated killing, but it is likely that recovery is at least in part due to decreased mutanobactin production by UA159. Enterococcal GelE may act directly on mutanobactin to degrade or inhibit it, or could process a specific component of the *E. faecalis* cell surface (such as a receptor or membrane protein) that is required for inhibition by mutanobactin. The latter could explain why the *S. aureus* extracellular proteases do not impact recovery from killing: if a *S. aureus* cell surface factor is required for mutanobactin-mediated killing then perhaps the *S. aureus* proteases do not process this factor in the same manner as GelE. It is also possible that mutanobactin induces transcriptional changes in *E. faecalis* that promote resistance over time. Future work will define the mechanism by which GelE contributes to recovery from mutanobactin-mediated killing.

In an effort to contextualize our results in a biologically relevant condition, we examined UA159 mutanobactin-mediated inhibition of OG1RF in the context of biofilms as *S. mutans* primarily exists within dental plaque biofilms on the surface of teeth. While planktonic UA159 production of mutanobactin inhibited OG1RF biofilm formation and killed pregrown OG1RF biofilms, UA159 biofilms did not exclude OG1RF from forming dual-species biofilms on Aclar (**Fig. 6E**) and inhibition of planktonic OG1RF by UA159 biofilms was slight (**Fig. S5C**). Although we pre-formed the UA159 biofilms and co-cultured them with OG1RF in BHI supplemented with 1% sucrose, which has been shown to reduce mutanobactin gene expression (78), planktonic UA159 killing of OG1RF was increased in 1% sucrose (**Fig. 4C**). This suggests that mutanobactin gene expression by planktonic UA159 may be upregulated in the presence of OG1RF while expression by biofilm-contained UA159 is downregulated. Alternatively, *E. faecalis* could be more susceptible to killing in the presence of sucrose due to increased acid production (78). Although *E. faecalis* viability is not affected at a culture pH as low as 4 (**Fig. S2B**), the effect of a lower pH and mutanobactin production could be additive, even with mutanobactin gene expression decreased in the presence of sucrose (78). Studies measuring mutanobactin gene expression under different conditions have shown that mutanobactin genes are upregulated in glucose and saliva (78, 79), supporting the idea that mutanobactin expression is context-dependent. The ability of OG1RF to form biofilms despite the presence of a UA159 biofilm could explain why *E. faecalis* is able to occupy the oral cavity in the presence of *S. mutans* since *S. mutans* is primarily found in biofilms rather than as planktonic cells. The mechanism underlying dual-species biofilm formation by *E. faecalis* and *S. mutans* is unclear, but we hypothesize that *S. mutans* and *E. faecalis* are spatially separated from each other within biofilms, thereby reducing cell-cell contact between the two species. This is consistent with a previous observation that the number of both *E. faecalis* and *S. mutans* cells within a dual-species biofilm are lower than the numbers of cells for either species in single-species biofilms (44). It is also possible that *S. mutans* does not produce mutanobactin within biofilms, which would be consistent with UA159 production of mutanobactin coinciding with exponential growth (57). Additional studies are needed to delineate the spatiotemporal dynamics of *S. mutans* and *E. faecalis* biofilm formation in the oral cavity, and how *E. faecalis* is able to be established as a primary endodontic pathogen.

Our work demonstrates the antibacterial effects of streptococcal natural products on Gram-positive bacteria, including ESKAPE pathogens. We identified two strategies *S. mutans* strains use to kill other bacteria, one mutanobactin-and contact-dependent and the other through the secretion of a different natural product independent of cell-cell contact. We show the potential utility of mutanobactin in preventing and treating biofilms formed by *E. faecalis*, a clinically important pathobiont that causes widespread infections in humans. While mutanobactin has previously been characterized, this work is the first demonstration of its antibacterial activity, highlighting the functional diversity of metabolites produced by microbes. The total synthesis of Streptococcal mutanobactin has been described (80, 81), providing a framework for the potential development of mutanobactin as a treatment for Gram-positive infections, including those resistant to other antibiotics. New and unknown streptococcal natural products may also be exploited for therapeutic control of Gram-positive infections. In this age of increased need for antibiotic discovery, our work underscores the benefits of identifying novel natural products from well-characterized species and investigating known natural products for potential antimicrobial activity.

## ACKNOWLEDGMENTS

This work was supported by R00AI151080 to JLEW. RYI was supported by an Institutional Research and Academic Career Development Award fellowship from NIH grant K12GM119955. We thank Bruno Lima, Mark Herzberg, Indranil Biswas, Anthony Baughn, and Alexander Horswill for sharing strains. We thank Bruno Lima and members of the Willett lab for providing feedback on the manuscript.

## AUTHOR CONTRIBUTIONS

RYI and JLEW designed the research; RYI and CWS performed experiments and analyzed data; RYI, CWS, and JLEW wrote the manuscript.

## MATERIALS AND METHODS

### Bacterial strains, plasmids, media, and growth conditions

Strains used in this study are listed in Table 1. Plasmids used in this study are listed in Table 2. Strains were grown in brain heart infusion (BHI) medium (per liter: 37 g Difco BHI [BD]) unless otherwise noted. OG1RF strains were grown in tryptic soy broth without added dextrose (TSB-D) medium (per liter: 27.5 g Bacto tryptic soy broth without dextrose [BD]) where noted. *E. coli* strains used for cloning were grown in Luria-Bertani (LB) medium (per liter: 25 g LB broth [Electron Microscopy Sciences]; or 10 g tryptone [Fisher Scientific], 5 g yeast extract [Bacto Difco or Electron Microscopy Sciences], 1 g glucose, 5 g NaCl) or BHI medium. When needed, antibiotics were added to media at the following concentrations: fusidic acid, 25 μg/mL for OG1RF; erythromycin, 10 μg/mL for OG1RF, 5 μg/mL for UA159; and tetracycline, 10 μg/mL for OG1RF and *E. coli*. When needed, the peptide pheromone cCF10 (50 ng/mL, Mimotopes) was added to media. When needed, 1% (w/v) sucrose was added to media. Growth medium was solidified using 1% agar when needed. For gelatin plates, 3% w/v gelatin was added to BHI agar. *Enterococcus* spp. and *Streptococcus* spp. were grown at 37°C in 5% CO2 without agitation unless otherwise noted. *Staphylococcus* spp*., K. pneumoniae, A. baumannii, P. aeruginosa,* and *E. aerogenes* were grown at 37°C with shaking unless otherwise noted.

**Table 2.** Plasmids used in this study.

| Plasmid | Description | Source or Reference(s) |
| --- | --- | --- |
| pMSP3535 | Shuttle vector with nisin-inducible promoter; pAMβ1 and ColE1 replicons, *nisRK*, *PnisA* (Erm<sup>R</sup>) | (106) |
| pJW76 | cCF10 pheromone-inducible plasmid for gene expression (Tet<sup>R</sup>) | (92) |
| pJW1016 | pJW76 harboring OG1RF *gelE* for cCF10 pheromone-inducible expression (Tet<sup>R</sup>) | This study |
| pCJK205 | Constitutive *lacZ* (Erm<sup>R</sup>) | (71) |

### DNA synthesis and sequencing

Primers used in this study are listed in Table 3 and were synthesized by ThermoFisher Scientific or Integrated DNA Technologies (Coralville, IA, USA). Full pGelE plasmid was confirmed by whole plasmid sequencing by Plasmidsaurus using Oxford Nanopore Technology with custom analysis and annotation. *S. mutans* UA159 erm^R^, Δ*mubD*, and Δ*mubR* strains were confirmed by whole genome sequencing performed by SeqCoast Genomics (Portsmouth, NH, USA) using Illumina short read sequencing with 150 bp paired end reads. PCR to amplify constructs for cloning and strain construction were performed using PfuUltra II or PfuUltra II Fusion HS DNA Polymerase (Agilent Technologies). Diagnostic PCR was performed using Taq or OneTaq DNA Polymerase (NEB).

**Table 3.**
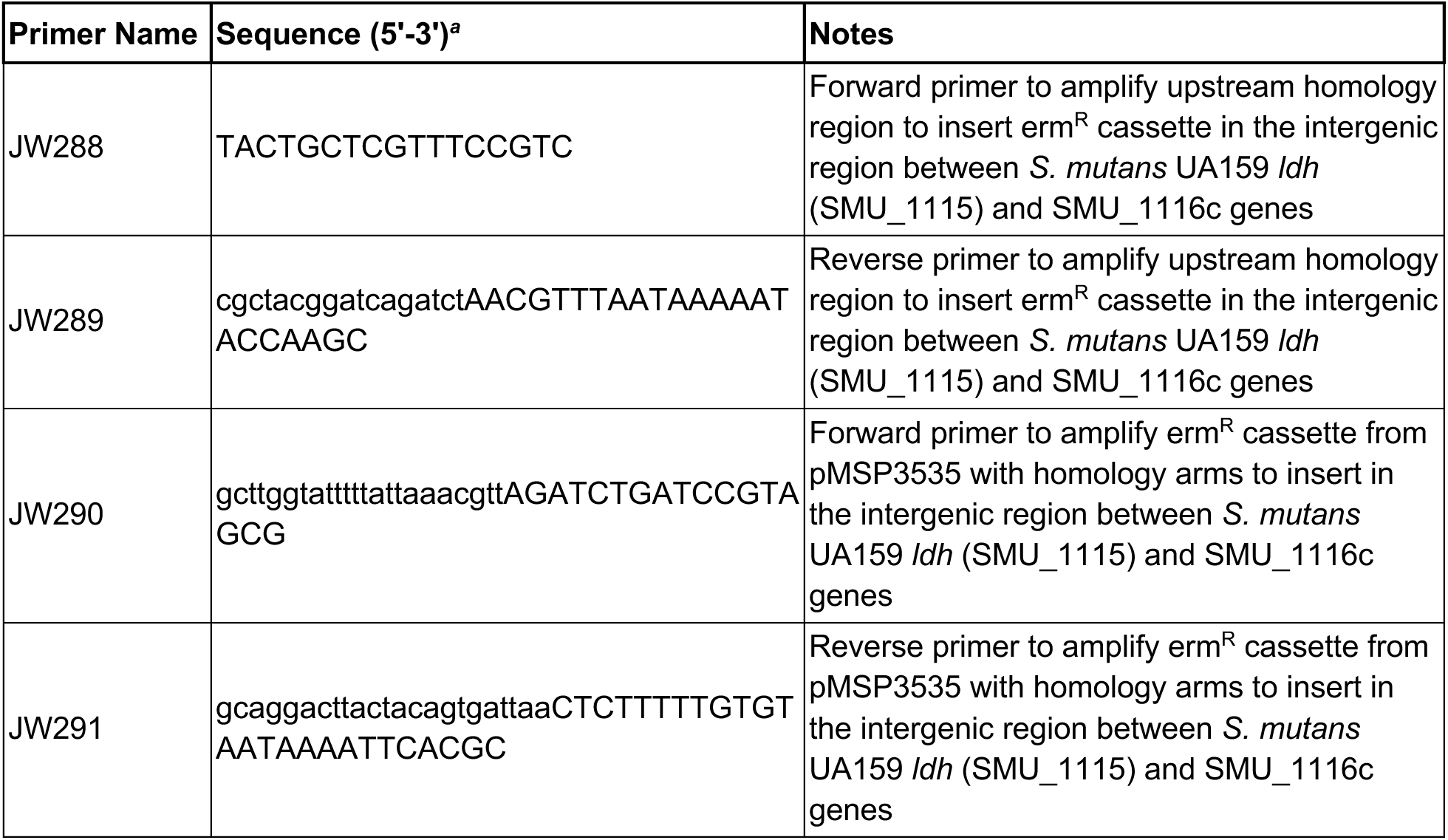
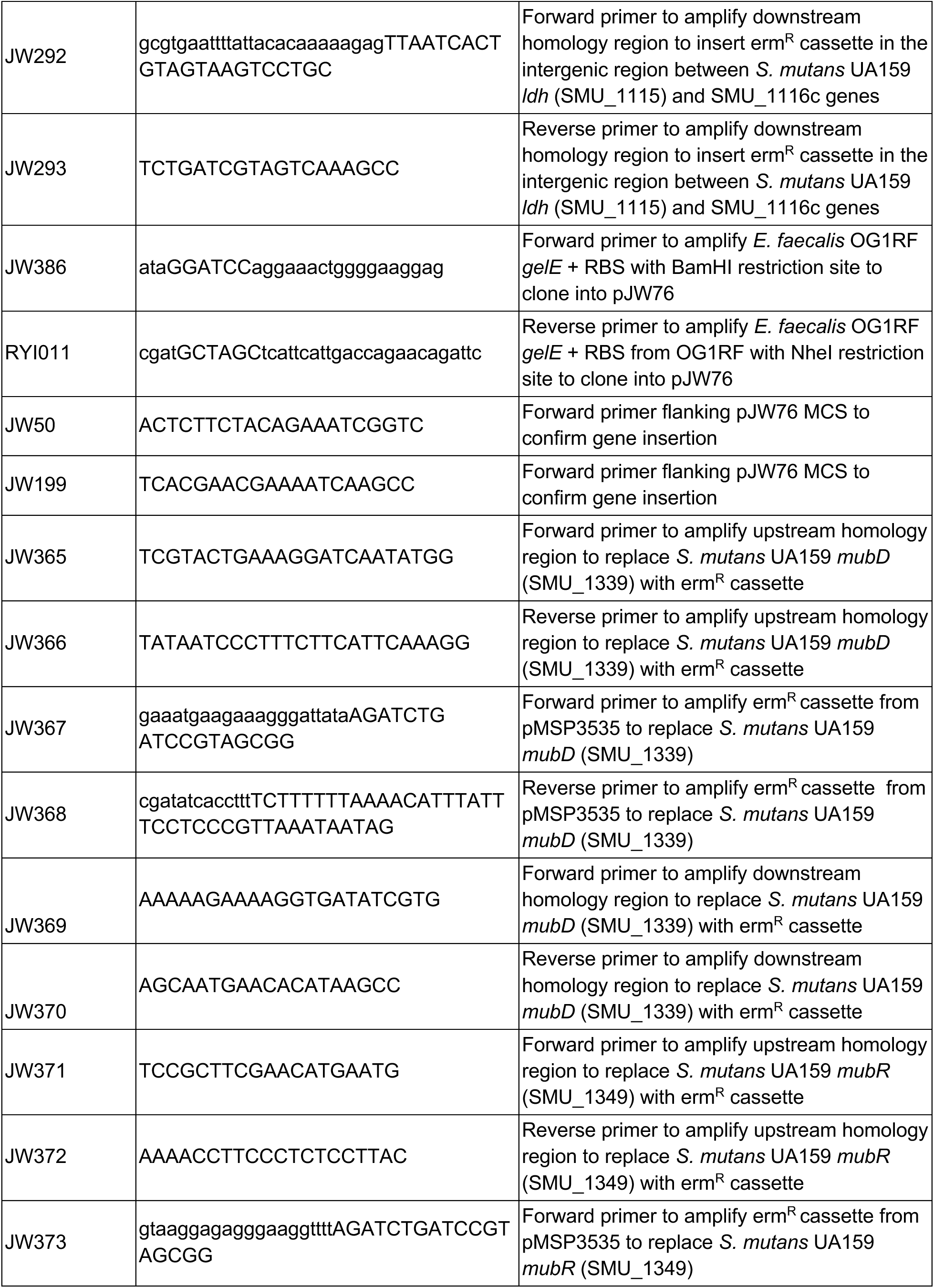
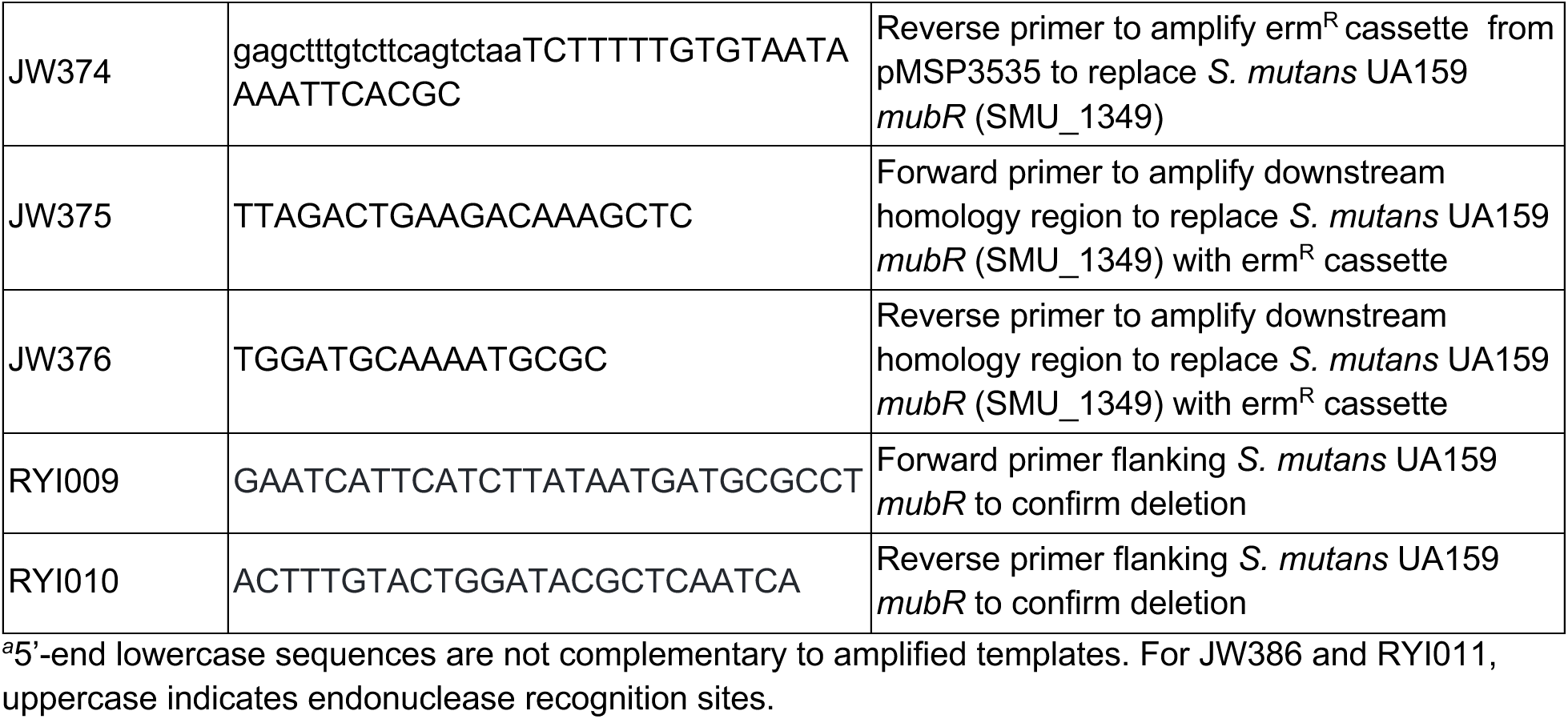
Primer list.

### Construction of *S. aureus* UA159 strains

Mutagenesis was performed according to an established protocol (82) to insert an erythromycin resistance cassette in the intergenic region between UA159 *ldh* (SMU_1115) and SMU_1116c, or to replace UA159 *mubD* or *mubR*. For the *ldh*/SMU_1116c IG::*erm* strain, ∼600-800 bp of the upstream and downstream regions flanking the *ldh*/SMU_1116c intergenic insertion site were amplified by PCR using primer pairs JW288/JW289 and JW292/JW293, respectively. For the Δ*mubD*::*erm* strain, ∼900-1000 bp of the upstream and downstream regions flanking *mubD* were amplified using primer pairs JW365/JW366 and JW369/JW370, respectively. For the Δ*mubR*::*erm* strain, ∼1000 bp of the upstream and downstream regions flanking *mubR* were amplified using primer pairs JW371/JW372 and JW375/JW376, respectively. For all mutants, the erm^R^ cassette from pMSP3535 was amplified using primers JW290 and JW291. For *ldh*/SMU_1116c IG::*erm* and Δ*mubD*::*erm*, the resulting PCR products were treated with DpnI and cleaned over a silica column (Epoch). The upstream, downstream, and erm^R^ PCR products for each strain were then spliced together by overlap extension PCR (OE-PCR) using the forward primer used to amplify the upstream region and the reverse primer used to amplify the downstream region. The resulting linear mutagenic DNA fragments were transformed into UA159 and transformants were selected using erythromycin. Strains were confirmed by PCR using primers flanking the deleted gene followed by whole genome sequencing.

### Construction of pJW1016 *gelE* expression plasmid

*E. faecalis* OG1RF *gelE* was PCR amplified from genomic DNA using primers JW386 and RYI011 that added flanking BamHI and NheI restriction sites. The amplified *gelE* and pJW76 were digested with BamHI and NheI (NEB), then digested *gelE* was ligated into digested pJW76 using T4 DNA ligase (NEB). The ligation reaction was transformed into chemically competent DH5α. Τransformants were selected using tetracycline and confirmed by PCR followed by whole plasmid sequencing. Electrocompetent OG1RF Δ*gelE* was transformed with pJW1016, transformants were selected using tetracycline, and strain was confirmed by PCR.

### Co-culture assay

Overnight cultures of target strains were diluted to 10^7^ CFU/mL or OD600 of 0.01 in BHI in a U-bottom 96-well plate. Overnight cultures of competing strains were inoculated with the target strain at the corresponding CFU/mL or OD600 for 1:1, 1:2, 1:5, or 1:10 co-cultures. Co-cultures were incubated at 37°C in 5% CO2. For each time point, samples of co-cultures were removed, diluted in potassium based phosphate buffered saline (KPBS; 10 mM, pH 7.4), and spotted on BHI agar to quantify CFU/mL of each strain in co-culture. For CFU enumeration, fusidic acid and/or incubation without 5% CO2 was used to select for OG1RF strains, incubation without 5% CO2 was used to select for all other target strains, and erythromycin was used to select for UA159 erm^R^, Δ*mubD,* and Δ*mubR* strains.

### Gelatinase assay

Overnight cultures of *E. faecalis* clinical isolates were spotted onto BHI agar supplemented with 3% (w/v) gelatin and incubated overnight (∼24 h) at 37°C. Plates were imaged using a ProteinSimple Cell Biosciences FlourChem FC3 System imager. The radius of the opaque gelatinase halo for each strain was measured from the edge of the colony to the edge of the gelatinase halo using FIJI (83).

### Chlorophenyl red-β-D-galactopyranoside (CPRG) cell membrane permeability assay

CPRG assays were performed according to previously described methods (70, 71). Overnight cultures of OG1RF pCJK205 grown in TSB-D-Erm^10^ were diluted to 10^7^ CFU/mL in TSB-D with CPRG (25 μg/mL) in a glass culture tube and incubated at 37°C with 5% CO2 for 24 h. For antibiotic CPRG assays, subinhibitory concentrations of daptomycin (8 μg/mL + 50 μg/mL CaCl2) and ciprofloxacin (0.5 μg/mL) were added to media. For co-culture CPRG assays, overnight cultures of UA159 strains grown in TSB-D-Erm^10^ were inoculated with OG1RF pCJK205 at 10^8^ CFU/mL. CPRG cleavage was measured by pelleting cells at 8000 x g for 2 min, transferring the supernatant to a flat-bottom 96-well plate, and measuring A570 using a BioTek Synergy H1 plate reader. Images of CPRG assay culture tubes were taken using a cell phone camera.

### Supernatant growth assay

Overnight cultures of strains were diluted to 10^8^ CFU/mL in BHI and incubated for 4 h at 37°C with 5% CO2. For co-culture supernatants, OG1RF was inoculated at 10^7^ CFU/mL with UA159 or ATCC 33535 at 10^8^ CFU/mL. Cell-free supernatants were obtained by removing cells via centrifugation at 8000 x g for 2 min followed by filter-sterilization of supernatants using a 0.45 μm filter. Overnight cultures of OG1RF were diluted to 10^7^ CFU/mL in a 50/50 mixture of BHI and cell-free supernatant and incubated for 4 h at 37°C with 5% CO2. To quantify OG1RF CFU/mL, cultures were diluted in KPBS (10 mM, pH 7.4), and spotted on BHI agar.

### Aclar biofilm assays

Overnight cultures of OG1RF or UA159 strains were diluted to 10^7^ CFU/mL or 10^8^ CFU/mL, respectively, in BHI (+1% sucrose for UA159) in a 48-well plate containing a sterile 5 mm diameter Aclar fluoropolymer disk (Electron Microscopy Sciences). For co-culture biofilm assays, overnight cultures of UA159 strains were inoculated with OG1RF at 10^7^ or 10^8^ CFU/mL and incubated at 37°C with 5% CO2. For pre-formed biofilm assays, OG1RF and UA159 biofilms were incubated at 37°C with 5% CO2 overnight, then Aclar disks were washed x3 in KPBS (10 mM, pH 7.4) before transferring to fresh BHI containing 10^7^ OG1RF or 10^8^ CFU/mL UA159 strains and incubated at 37°C with 5% CO2. At 4 and 24 h, planktonic cells were removed, diluted in KPBS, and spotted on BHI agar to quantify CFU/mL of each strain. To quantify CFUs of Aclar biofilm cells at 4 and 24 h, Aclar disks were removed, washed x3 in KPBS, transferred to a 1.5 mL microcentrifuge tube containing 1 mL KPBS, vortexed for 5 min at 2500 rpm using a multi-tube vortexer, and then cells were diluted in KPBS and spotted on BHI agar. For CFU enumeration, fusidic acid and incubation without 5% CO2 was used to select for OG1RF strains and erythromycin was used to select for UA159 strains.

### Whole genome sequencing and genome analysis

Whole genome sequencing of *S. mutans* strains UA159B, Ingbritt, and ATCC 33535 were performed by SeqCoast Genomics (Portsmouth, NH, USA) using Illumina short read sequencing with 150 bp paired end reads. The UA159B genome sequence was compared to the available UA159 genome sequence (NC_004350.2) using breseq (84). Acquired and available *S. mutans* genomes were queried for *mubR* and *mubD* homologs using NCBI BLAST. The *mubR* homolog in ATCC 25175 (taxonomy ID 1257041) is locus tag D820_RS03580/D820_06113 on contig008 of reference sequence NZ_AOCB01000008.1. The *mubR* and *mubD* genes in UA159 and UA140 were identified from published work (57).

### Data analysis

CFU/mL data were log10 transformed before performing statistical analysis. Limit of detection (LOD) for CFU/mL was determined by calculating CFU/mL based on the smallest number of colonies (1) that could be detected at the lowest dilution (10^-1^) plated. GraphPad Prism was used to generate graphs and conduct statistical analyses. Graphs and figures were assembled and refined in Adobe Illustrator.

## DATA, MATERIALS, AND SOFTWARE AVAILABILITY

Genome sequencing files for *S. mutans* isolates UA159, UA159B, Ingbritt, and ATCC 33535 have been deposited with NCBI Genome under BioProject ID PRJNA974474.

## FIGURE LEGENDS

**Fig. S1.**
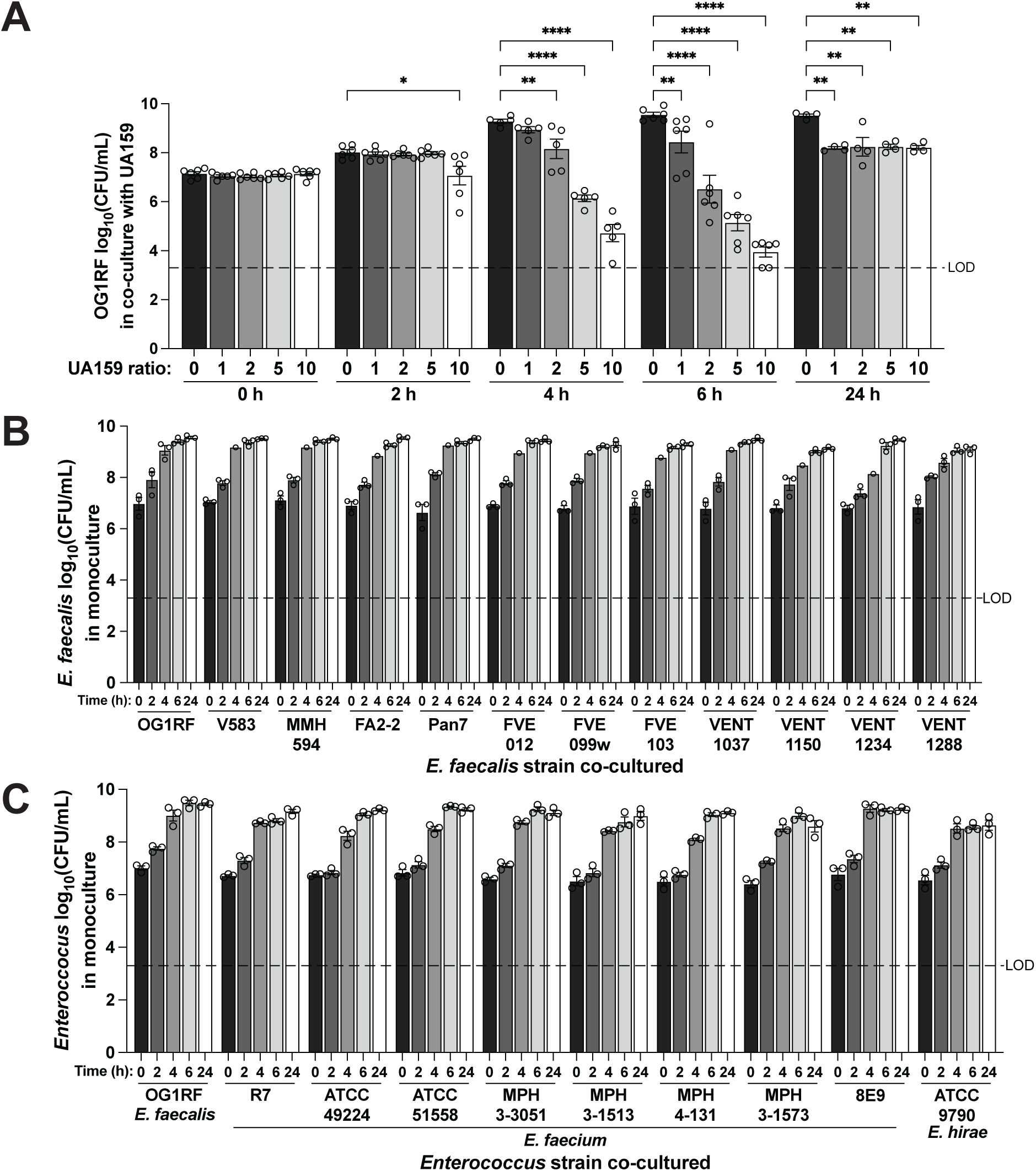
*S. mutans* UA159 kills *E. faecalis* OG1RF in a dose-dependent manner. **A.** *E. faecalis* OG1RF CFU/mL in co-culture with indicated *S. mutans* UA159 erm^R^ at indicated inoculum ratio of UA159 erm^R^ to 1 part OG1RF in BHI at 0, 2, 4, 6, and 24 h. For each culture, n = 2 technical replicates for each of n = 4-6 biological replicates. Two-way ANOVA was used for statistical analysis (ns, not significant; *, p = 0.0149; **, p ≤ 0.0075; ****, p < 0.0001). **B.** CFU/mL of indicated *E. faecalis* clinical isolates in BHI monoculture at 0, 2, 4, 6, and 24 h. For each culture, n = 2 technical replicates for each of n = 1-3 biological replicates. **C.** CFU/mL of indicated *Enterococcus* spp. strains in BHI monoculture at 0, 2, 4, 6, and 24 h. For each culture, n = 2 technical replicates for each of n = 3 biological replicates. For A-C, each dot represents the mean of technical replicates, bars represent the mean of biological replicates, error bars represent standard errors of the mean, dashed lines indicate limit of detection (LOD).

**Fig. S2.**
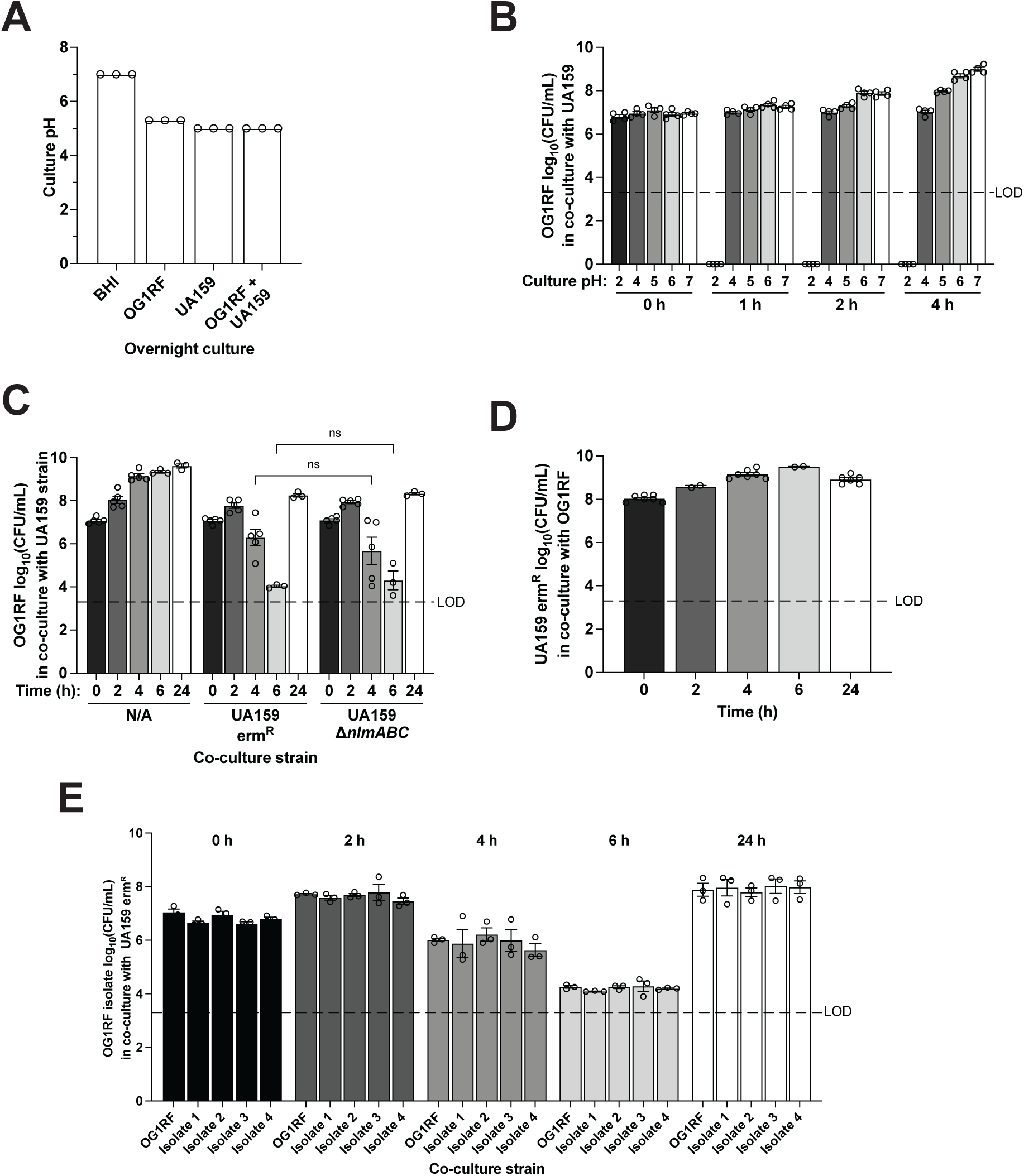
OG1RF viability is not impacted by culture pH change by UA159 or by UA159 mutacin production, OG1RF does not acquire lasting resistance to killing. **A.** pH of indicated overnight monoculture or co-culture. For each culture, n = 3 biological replicates. B. *E. faecalis* OG1RF CFU/mL cultured in BHI at indicated pH at 0, 1, 2, and 4 h. For each culture, n = 2 technical replicates for each of n = 4 biological replicates. C. *E. faecalis* OG1RF CFU/mL in co-culture with indicated *S. mutans* UA159 strain in BHI at 0, 2, 4, 6, and 24 h. For each culture, n = 2 technical replicates for each of n = 3-5 biological replicates. Two-way ANOVA was used for statistical analysis (ns, not significant). D. *S. mutans* UA159 erm^R^ CFU/mL in 10:1 co-culture with *E. faecalis* OG1RF in BHI at 0, 2, 4, 6, and 24 h. For each culture, n = 2 technical replicates for each of n = 2-7 biological replicates. **E.** CFU/mL of *E. faecalis* OG1RF and OG1RF 24 h co-culture isolates in 1:10 co-culture with *S. mutans* UA159 erm^R^ in BHI at 0, 2, 4, 6, and 24 h. For each culture, n = 2 technical replicates for each of n = 3 biological replicates. For A-E, each dot represents the mean of technical replicates, bars represent the mean of biological replicates, error bars represent standard errors of the mean, dashed lines indicate limit of detection (LOD), and data points at y = 0 indicate no CFUs were detected.

**Fig. S3.**
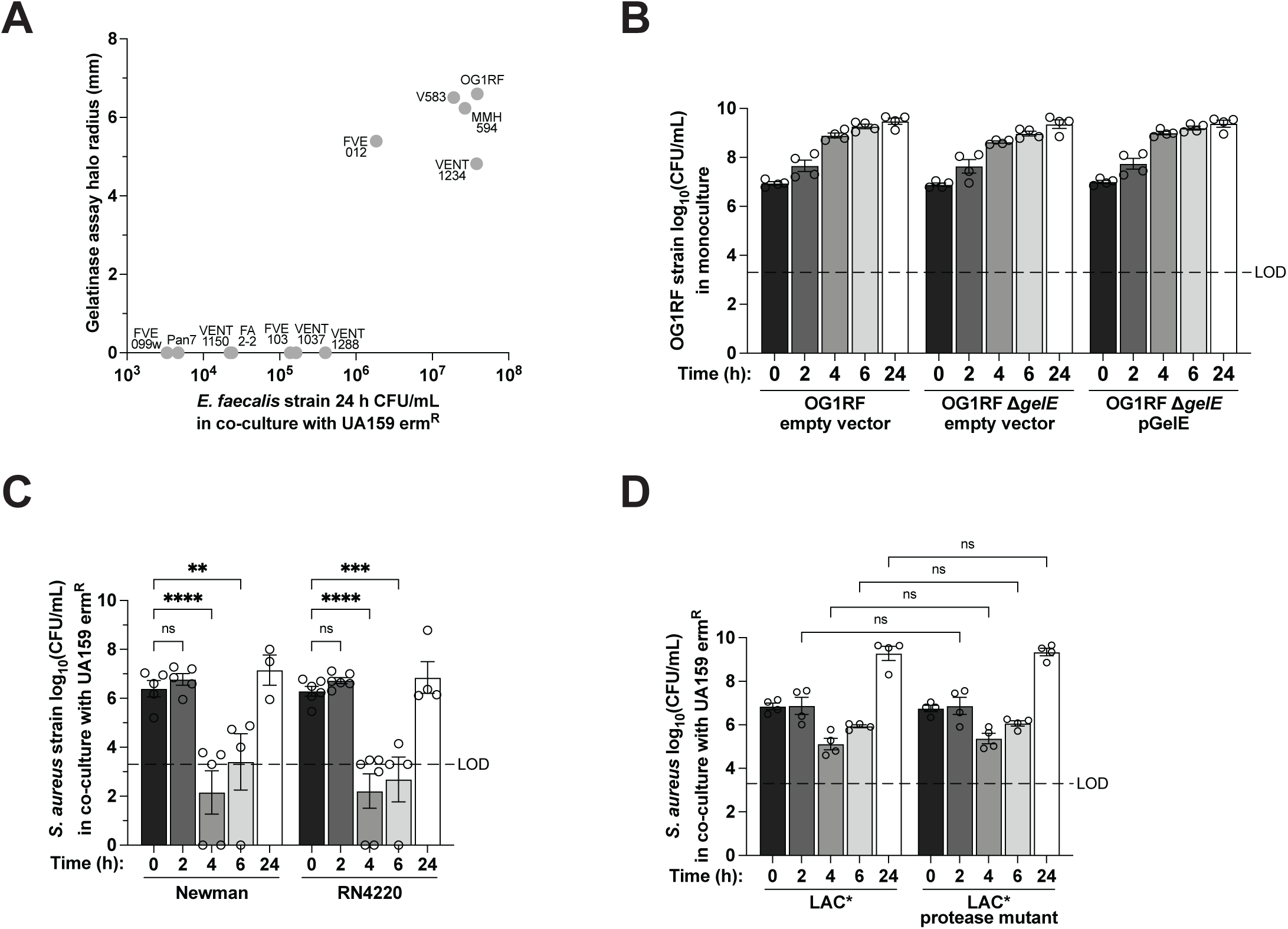
*E. faecalis* gelatinase activity is required for resistance to killing by *S. mutans* UA159, *S. aureus* extracellular proteases are not required for resistance. A. Correlation between *E. faecalis* strain 24 h CFU/mL in co-culture with *S. mutans* UA159 erm^R^ and *E. faecalis* strain gelatinase activity. Each dot represents the mean of biological replicates for a strain of *E. faecalis*. **B.** CFU/mL of indicated *S. aureus* strain monoculture in BHI at 0, 2, 4, 6, and 24 h. For each culture, n = 2 technical replicates for each of n = 4-6 biological replicates. **C.** CFU/mL of indicated *S. aureus* strain in 1:10 co-culture with *S. mutans* UA159 erm^R^ in BHI at 0, 2, 4, 6, and 24 h. For each culture, n = 2 technical replicates for each of n = 4-6 biological replicates. **D.** CFU/mL of indicated *S. aureus* LAC* strains in 1:10 co-culture with *S. mutans* UA159 erm^R^ in BHI at 0, 2, 4, 6, and 24 h. For each culture, n = 2 technical replicates for each of n = 4 biological replicates. For B-D, each dot represents the mean of technical replicates, bars represent the mean of biological replicates, error bars represent standard errors of the mean, dashed lines indicate limit of detection (LOD), and data points at y = 0 indicate no CFUs were detected. For C and D, two-way ANOVA was used for statistical analysis (ns, not significant; **, p = 0.0074; ***, p = 0.0007; ****, p < 0.0001).

**Fig. S4.**
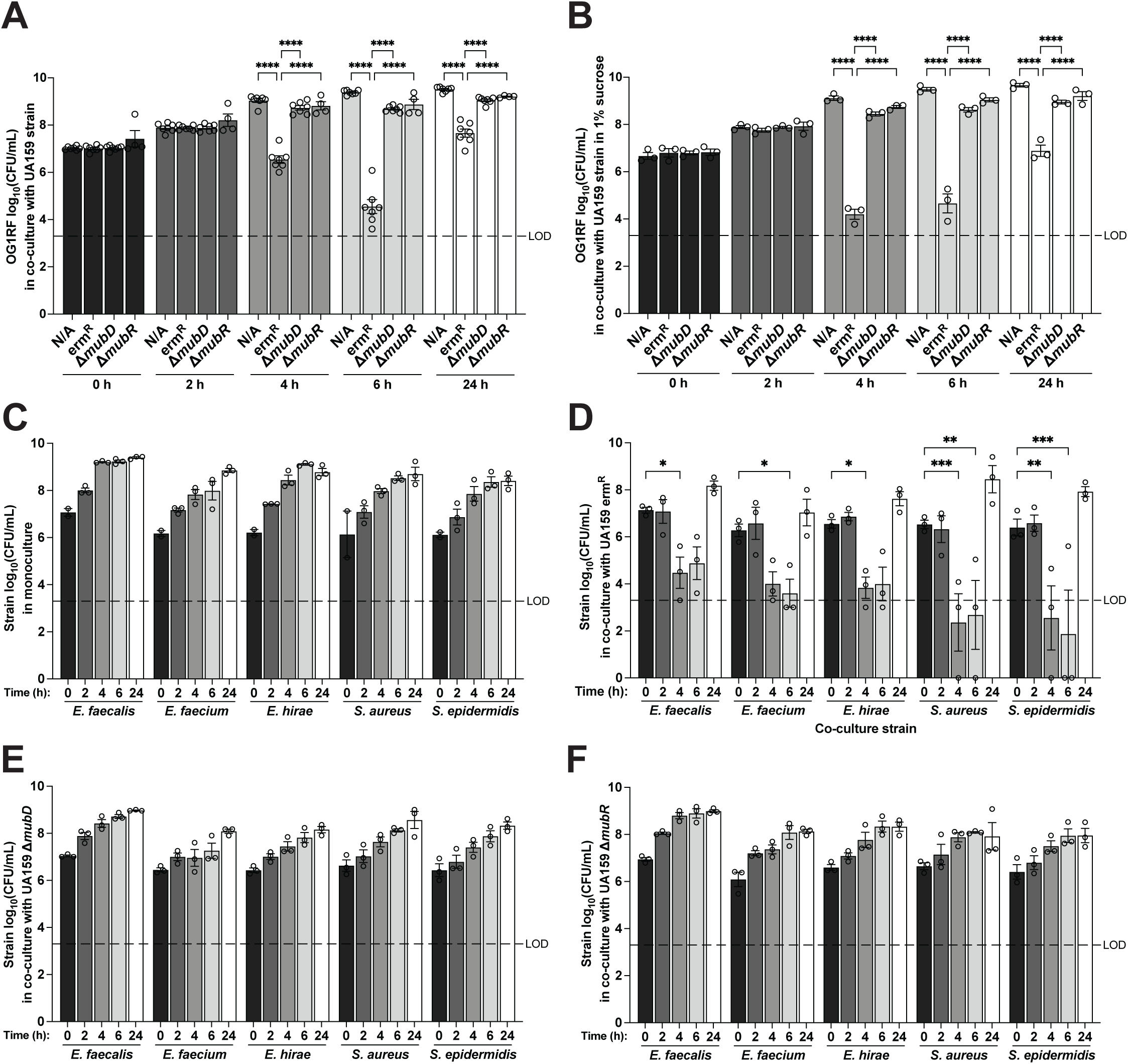
*S. mutans* UA159 mutanobactin genes are required to kill Gram-positive pathogens in co-culture. **A.** *E. faecalis* OG1RF CFU/mL in 1:10 co-culture with indicated *S. mutans* UA159 strain in BHI at 0, 2, 4, 6, and 24 h. For each culture, n = 2 technical replicates for each of n = 4-7 biological replicates. **B.** *E. faecalis* OG1RF CFU/mL in 1:10 co-culture with indicated *S. mutans* UA159 strain in BHI + 1% sucrose at 0, 2, 4, 6, and 24 h. For each culture, n = 2 technical replicates for each of n = 3 biological replicates. **C.** CFU/mL of indicated strain in monoculture in BHI at 0, 2, 4, 6, and 24 h. For each culture, n = 2 technical replicates for each of n = 2-3 biological replicates. **D.** CFU/mL of indicated strain in 1:10 co-culture with *S. mutans* UA159 erm^R^ in BHI at 0, 2, 4, 6, and 24 h. For each culture, n = 2 technical replicates for each of n = 3 biological replicates. **E.** CFU/mL of indicated strain in 1:10 co-culture with *S. mutans* UA159 Δ*mubD* in BHI at 0, 2, 4, 6, and 24 h. **F.** CFU/mL of indicated strain in 1:10 co-culture with *S. mutans* UA159 Δ*mubR* in BHI at 0, 2, 4, 6, and 24 h. For each culture, n = 2 technical replicates for each of n = 3 biological replicates. For A, B, and D, two-way ANOVA was used for statistical analysis (*, p < 0.05; **, p = 0.002; ***, p ≤ 0.0008; ****, p < 0.0001).

**Fig. S5.**
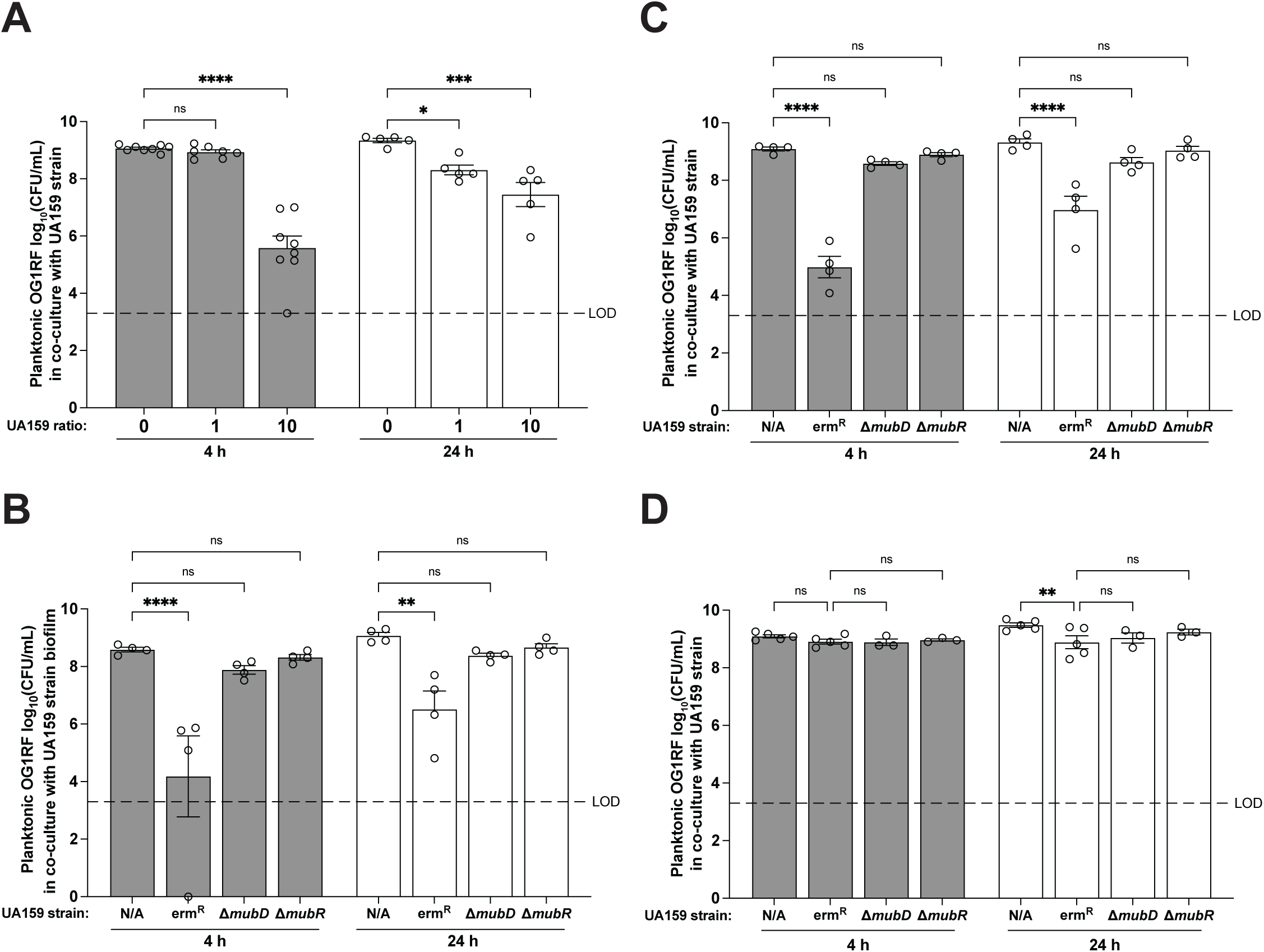
Mutanobactin production by *S. mutans* UA159 kills *E. faecalis* OG1RF planktonic cells in the presence of an Aclar disk. **A.** Planktonic *E. faecalis* OG1RF CFU/mL in co-culture with *S. mutans* UA159 erm^R^ at indicated inoculum ratio of UA159 erm^R^ to 1 part OG1RF in BHI at 4 and 24 h. For each co-culture, n = 2 technical replicates for each of n = 7-8 biological replicates. **B.** Planktonic *E. faecalis* OG1RF CFU/mL in 1:10 co-culture with indicated *S. mutans* UA159 strain in BHI at 4 and 24 h. For each co-culture, n = 2 technical replicates for each of n = 3-4 biological replicates. **C.** Planktonic *E. faecalis* OG1RF CFU/mL of OG1RF pre-formed Aclar biofilm in co-culture with indicated *S. mutans* UA159 strain in BHI at 4 and 24 h. For each co-culture, n = 2 technical replicates for each of n = 4 biological replicates. **D.** Planktonic *E. faecalis* OG1RF CFU/mL in co-culture with pre-formed Aclar biofilm of indicated *S. mutans* UA159 strain in BHI at 4 and 24 h. For each co-culture, n = 2 technical replicates for each of n = 3-5 biological replicates. For A-D, two-way ANOVA was used for statistical analysis (ns, not significant; *, p = 0.0374; **, p < 0.003; ***, p = 0.0002; ****, p < 0.0001), each dot represents the mean of technical replicates, bars represent the mean of biological replicates, error bars represent standard errors of the mean, dashed lines indicate limit of detection (LOD), and data points at y = 0 indicate no CFUs were detected.

## REFERENCES

1. Rice LB. 2008. Federal funding for the study of antimicrobial resistance in nosocomial pathogens: no ESKAPE. J Infect Dis 197:1079–1081.

2. Miller WR, Arias CA. 2024. ESKAPE pathogens: antimicrobial resistance, epidemiology, clinical impact and therapeutics. Nat Rev Microbiol 22:598–616.

3. CDC. 2025. CDC Partners Estimate Healthcare Cost of Antimicrobial-resistant Infections. Antimicrobial Resistance. https://www.cdc.gov/antimicrobial-resistance/stories/partner-estimates.html. Retrieved 24 June 2025.

4. Flores-Mireles AL, Walker JN, Potretzke A, Schreiber HL IV, Pinkner JS, Bauman TM, Park AM, Desai A, Hultgren SJ, Caparon MG. 2016. Antibody-based therapy for enterococcal catheter-associated urinary tract infections. MBio 7:e01653–16.

5. Guzman Prieto AM, van Schaik W, Rogers MRC, Coque TM, Baquero F, Corander J, Willems RJL. 2016. Global emergence and dissemination of enterococci as nosocomial pathogens: Attack of the clones? Front Microbiol 7:788.

6. Barnes AMT, Dale JL, Chen Y, Manias DA, Greenwood Quaintance KE, Karau MK, Kashyap PC, Patel R, Wells CL, Dunny GM. 2017. *Enterococcus faecalis* readily colonizes the entire gastrointestinal tract and forms biofilms in a germ-free mouse model. Virulence 8:282–296.

7. Tien BYQ, Goh HMS, Chong KKL, Bhaduri-Tagore S, Holec S, Dress R, Ginhoux F, Ingersoll MA, Williams RBH, Kline KA. 2017. *Enterococcus faecalis* promotes innate immune suppression and polymicrobial catheter-associated urinary tract infection. Infect Immun 85:e00378–17.

8. Madsen KT, Skov MN, Gill S, Kemp M. 2017. Virulence factors associated with *Enterococcus faecalis* infective endocarditis: A mini review. Open Microbiol J 11:1–11.

9. Fiore E, Van Tyne D, Gilmore MS. 2019. Pathogenicity of enterococci. Microbiol Spectr 7:GPP3–0053–2018.

10. Zhang Y, Du M, Chang Y, Chen L-A, Zhang Q. 2017. Incidence, clinical characteristics, and outcomes of nosocomial *Enterococcus* spp. bloodstream infections in a tertiary-care hospital in Beijing, China: a four-year retrospective study. Antimicrob Resist Infect Control 6:73.

11. Arias CA, Murray BE. 2012. The rise of the *Enterococcus*: beyond vancomycin resistance. Nat Rev Microbiol 10:266–278.

12. Zirakzadeh A, Patel R. 2006. Vancomycin-resistant enterococci: colonization, infection, detection, and treatment. Mayo Clin Proc 81:529–536.

13. Sahm DF, Kissinger J, Gilmore MS, Murray PR, Mulder R, Solliday J, Clarke B. 1989. *In vitro* susceptibility studies of vancomycin-resistant *Enterococcus faecalis*. Antimicrobial agents and chemotherapy 33:1588–1591.

14. Bell JM, Paton JC, Turnidge J. 1998. Emergence of vancomycin-resistant enterococci in Australia: phenotypic and genotypic characteristics of isolates. J Clin Microbiol 36:2187– 2190.

15. Shlaes DM, Bouvet A, Devine C, Shlaes JH, al-Obeid S, Williamson R. 1989. Inducible, transferable resistance to vancomycin in *Enterococcus faecalis* A256. Antimicrob Agents Chemother 33:198–203.

16. Shankar N, Baghdayan AS, Gilmore MS. 2002. Modulation of virulence within a pathogenicity island in vancomycin-resistant *Enterococcus faecalis*. Nature 417:746–750.

17. Paulsen IT, Banerjei L, Myers GSA, Nelson KE, Seshadri R, Read TD, Fouts DE, Eisen JA, Gill SR, Heidelberg JF, Tettelin H, Dodson RJ, Umayam L, Brinkac L, Beanan M, Daugherty S, DeBoy RT, Durkin S, Kolonay J, Madupu R, Nelson W, Vamathevan J, Tran B, Upton J, Hansen T, Shetty J, Khouri H, Utterback T, Radune D, Ketchum KA, Dougherty BA, Fraser CM. 2003. Role of mobile DNA in the evolution of vancomycin-resistant *Enterococcus faecalis*. Science 299:2071–2074.

18. Sharifi Y, Hasani A, Ghotaslou R, Varshochi M, Hasani A, Aghazadeh M, Milani M. 2012. Survey of Virulence Determinants among Vancomycin Resistant *Enterococcus faecalis* and *Enterococcus faecium* Isolated from Clinical Specimens of Hospitalized Patients of North west of Iran. Open Microbiol J 6:34–39.

19. Edelsberg J, Weycker D, Barron R, Li X, Wu H, Oster G, Badre S, Langeberg WJ, Weber DJ. 2014. Prevalence of antibiotic resistance in US hospitals. Diagn Microbiol Infect Dis 78:255–262.

20. Ubeda C, Taur Y, Jenq RR, Equinda MJ, Son T, Samstein M, Viale A, Socci ND, van den Brink MRM, Kamboj M, Pamer EG. 2010. Vancomycin-resistant *Enterococcus* domination of intestinal microbiota is enabled by antibiotic treatment in mice and precedes bloodstream invasion in humans. J Clin Invest 120:4332–4341.

21. Arias CA, Panesso D, McGrath DM, Qin X, Mojica MF, Miller C, Diaz L, Tran TT, Rincon S, Barbu EM, Reyes J, Roh JH, Lobos E, Sodergren E, Pasqualini R, Arap W, Quinn JP, Shamoo Y, Murray BE, Weinstock GM. 2011. Genetic basis for *in vivo* daptomycin resistance in enterococci. N Engl J Med 365:892–900.

22. Liu Y, Wang Y, Wu C, Shen Z, Schwarz S, Du X-D, Dai L, Zhang W, Zhang Q, Shen J. 2012. First report of the multidrug resistance gene *cfr* in *Enterococcus faecalis* of animal origin. Antimicrob Agents Chemother 56:1650–1654.

23. Patel SN, Memari N, Shahinas D, Toye B, Jamieson FB, Farrell DJ. 2013. Linezolid resistance in *Enterococcus faecium* isolated in Ontario, Canada. Diagn Microbiol Infect Dis 77:350–353.

24. Souto R, Colombo APV. 2008. Prevalence of *Enterococcus faecalis* in subgingival biofilm and saliva of subjects with chronic periodontal infection. Arch Oral Biol 53:155–160.

25. Wang Q-Q, Zhang C-F, Chu C-H, Zhu X-F. 2012. Prevalence of *Enterococcus faecalis* in saliva and filled root canals of teeth associated with apical periodontitis. Int J Oral Sci 4:19– 23.

26. Colombo APV, Teles RP, Torres MC, Souto R, Rosalém WJ, Mendes MCS, Uzeda M. 2002. Subgingival microbiota of Brazilian subjects with untreated chronic periodontitis. J Periodontol 73:360–369.

27. Gold OG, Jordan HV, van Houte J. 1975. The prevalence of enterococci in the human mouth and their pathogenicity in animal models. Arch Oral Biol 20:473–477.

28. Rams TE, Feik D, Young V, Hammond BF, Slots J. 1992. Enterococci in human periodontitis. Oral Microbiol Immunol 7:249–252.

29. Gaeta C, Marruganti C, Ali IAA, Fabbro A, Pinzauti D, Santoro F, Neelakantan P, Pozzi G, Grandini S. 2023. The presence of *Enterococcus faecalis* in saliva as a risk factor for endodontic infection. Front Cell Infect Microbiol 13:1061645.

30. Sedgley CM, Lennan SL, Clewell DB. 2004. Prevalence, phenotype and genotype of oral enterococci. Oral Microbiol Immunol 19:95–101.

31. Sedgley CM, Nagel AC, Shelburne CE, Clewell DB, Appelbe O, Molander A. 2005. Quantitative real-time PCR detection of oral *Enterococcus faecalis* in humans. Arch Oral Biol 50:575–583.

32. Sedgley C, Buck G, Appelbe O. 2006. Prevalence of *Enterococcus faecalis* at multiple oral sites in endodontic patients using culture and PCR. J Endod 32:104–109.

33. Souto R, Andrade AFB de, Uzeda M, Colombo APV. 2006. Prevalence of “non-oral” pathogenic bacteria in subgingival biofilm of subjects with chronic periodontitis. Braz J Microbiol 37:208–215.

34. Anderson AC, Jonas D, Huber I, Karygianni L, Wölber J, Hellwig E, Arweiler N, Vach K, Wittmer A, Al-Ahmad A. 2015. *Enterococcus faecalis* from food, clinical specimens, and oral sites: Prevalence of virulence factors in association with biofilm formation. Front Microbiol 6:1534.

35. Komiyama EY, Lepesqueur LSS, Yassuda CG, Samaranayake LP, Parahitiyawa NB, Balducci I, Koga-Ito CY. 2016. *Enterococcus* species in the oral cavity: Prevalence, virulence factors and antimicrobial susceptibility. PLoS One 11:e0163001.

36. Bakshi U, Sarkar M, Paul S, Dutta C. 2016. Assessment of virulence potential of uncharacterized *Enterococcus faecalis* strains using pan genomic approach - Identification of pathogen-specific and habitat-specific genes. Sci Rep 6:38648.

37. Stuart CH, Schwartz SA, Beeson TJ, Owatz CB. 2006. *Enterococcus faecalis*: its role in root canal treatment failure and current concepts in retreatment. J Endod 32:93–98.

38. Gaca AO, Lemos JA. 2019. Adaptation to adversity: The intermingling of stress tolerance and pathogenesis in enterococci. Microbiol Mol Biol Rev 83:e00008–19.

39. Colombo AV, Barbosa GM, Higashi D, di Micheli G, Rodrigues PH, Simionato MRL. 2013. Quantitative detection of *Staphylococcus aureus*, *Enterococcus faecalis* and *Pseudomonas aeruginosa* in human oral epithelial cells from subjects with periodontitis and periodontal health. J Med Microbiol 62:1592–1600.

40. McLean AR, Torres-Morales J, Dewhirst FE, Borisy GG, Mark Welch JL. 2022. Site-tropism of streptococci in the oral microbiome. Mol Oral Microbiol 37:229–243.

41. Mark Welch JL, Dewhirst FE, Borisy GG. 2019. Biogeography of the oral microbiome: The site-specialist hypothesis. Annu Rev Microbiol 73:335–358.

42. Lima AR, Ganguly T, Walker AR, Acosta N, Francisco PA, Pileggi R, Lemos JA, Gomes BPFA, Abranches J. 2020. Phenotypic and genotypic characterization of *Streptococcus mutans* strains isolated from endodontic infections. J Endod 46:1876–1883.

43. Lima AR, Herrera DR, Francisco PA, Pereira AC, Lemos J, Abranches J, Gomes BPFA. 2021. Detection of *Streptococcus mutans* in symptomatic and asymptomatic infected root canals. Clin Oral Investig 25:3535–3542.

44. Li X, Hoogenkamp MA, Ling J, Crielaard W, Deng DM. 2014. Diversity of *Streptococcus mutans* strains in bacterial interspecies interactions: Strain diversity and bacterial interspecies interaction. J Basic Microbiol 54:97–103.

45. Lemos JA, Palmer SR, Zeng L, Wen ZT, Kajfasz JK, Freires IA, Abranches J, Brady LJ. 2019. The biology of *Streptococcus mutans*. Microbiol Spectr 7:GPP3–0051–2018.

46. Hale JDF, Ting Y-T, Jack RW, Tagg JR, Heng NCK. 2005. Bacteriocin (mutacin) production by *Streptococcus mutans* genome sequence reference strain UA159: Elucidation of the antimicrobial repertoire by genetic dissection. Appl Environ Microbiol 71:7613–7617.

47. Hossain MS, Biswas I. 2011. Mutacins from *Streptococcus mutans* UA159 are active against multiple streptococcal species. Appl Environ Microbiol 77:2428–2434.

48. Su YA, Sulavik MC, He P, Makinen KK, Makinen PL, Fiedler S, Wirth R, Clewell DB. 1991. Nucleotide sequence of the gelatinase gene (*gelE*) from *Enterococcus faecalis* subsp. *liquefaciens*. Infect Immun 59:415–420.

49. Hancock LE, Perego M. 2004. Systematic inactivation and phenotypic characterization of two-component signal transduction systems of *Enterococcus faecalis* V583. J Bacteriol 186:7951–7958.

50. Qin X, Singh KV, Weinstock GM, Murray BE. 2000. Effects of *Enterococcus faecalis fsr* Genes on Production of Gelatinase and a Serine Protease and Virulence. Infect Immun 68:2579–2586.

51. Willett JLE, Dunny GM. 2025. Insights into ecology, pathogenesis, and biofilm formation of *Enterococcus faecalis* from functional genomics. Microbiol Mol Biol Rev 89:e0008123.

52. Murray BE, Singh KV, Ross RP, Heath JD, Dunny GM, Weinstock GM. 1993. Generation of restriction map of *Enterococcus faecalis* OG1 and investigation of growth requirements and regions encoding biosynthetic function. J Bacteriol 175:5216–5223.

53. Nakayama J, Cao Y, Horii T, Sakuda S, Akkermans AD, de Vos WM, Nagasawa H. 2001. Gelatinase biosynthesis-activating pheromone: a peptide lactone that mediates a quorum sensing in *Enterococcus faecalis*. Mol Microbiol 41:145–154.

54. Dubin G. 2002. Extracellular Proteases of *Staphylococcus* spp. Biol Chem 383:1075–1086.

55. Shaw L, Golonka E, Potempa J, Foster SJ. 2004. The role and regulation of the extracellular proteases of *Staphylococcus aureus*. Microbiology 150:217–228.

56. Joyner PM, Liu J, Zhang Z, Merritt J, Qi F, Cichewicz RH. 2010. Mutanobactin A from the human oral pathogen *Streptococcus mutans* is a cross-kingdom regulator of the yeast-mycelium transition. Org Biomol Chem 8:5486–5489.

57. Wu C, Cichewicz R, Li Y, Liu J, Roe B, Ferretti J, Merritt J, Qi F. 2010. Genomic island TnSmu2 of *Streptococcus mutans* harbors a nonribosomal peptide synthetase-polyketide synthase gene cluster responsible for the biosynthesis of pigments involved in oxygen and H_2_O_2_ tolerance. Appl Environ Microbiol 76:5815–5826.

58. Zvanych R, Lukenda N, Li X, Kim JJ, Tharmarajah S, Magarvey NA. 2015. Systems biosynthesis of secondary metabolic pathways within the oral human microbiome member *Streptococcus mutans*. Mol Biosyst 11:97–104.

59. Rainey K, Wilson L, Barnes S, Wu H. 2019. Quantitative proteomics uncovers the interaction between a virulence factor and mutanobactin synthetases in *Streptococcus mutans*. mSphere 4:e00429–19.

60. Wang X, Du L, You J, King JB, Cichewicz RH. 2012. Fungal biofilm inhibitors from a human oral microbiome-derived bacterium. Org Biomol Chem 10:2044–2050.

61. Paes Leme AF, Koo H, Bellato CM, Bedi G, Cury JA. 2006. The role of sucrose in cariogenic dental biofilm formation--new insight. J Dent Res 85:878–887.

62. Bowen WH. 2002. Do We Need to be Concerned about Dental Caries in the Coming Millennium? Crit Rev Oral Biol Med 13:126–131.

63. Marsh PD. 2003. Are dental diseases examples of ecological catastrophes? Microbiology 149:279–294.

64. Cai J-N, Jung J-E, Lee M-H, Choi H-M, Jeon J-G. 2018. Sucrose challenges to *Streptococcus mutans* biofilms and the curve fitting for the biofilm changes. FEMS Microbiol Ecol 94:fiy091.

65. Duarte S, Klein MI, Aires CP, Cury JA, Bowen WH, Koo H. 2008. Influences of starch and sucrose on *Streptococcus mutans* biofilms. Oral Microbiol Immunol 23:206–212.

66. Bowen WH, Koo H. 2011. Biology of *Streptococcus mutans*-derived glucosyltransferases: role in extracellular matrix formation of cariogenic biofilms. Caries Res 45:69–86.

67. Debono M, Abbott BJ, Molloy R, Fukuda D, Hunt A, Daupert V, Counter FT, Ott J, Carrell C, Howard L, Boeck LAVD, Hamill R. 1988. Enzymatic and chemical modifications of lipopeptide antibiotic A21978C: the synthesis and evaluation of daptomycin (LY146032). J Antibiot (Tokyo) 41:1093–1105.

68. Silverman JA, Perlmutter NG, Shapiro HM. 2003. Correlation of daptomycin bactericidal activity and membrane depolarization in *Staphylococcus aureus*. Antimicrob Agents Chemother 47:2538–2544.

69. Muraih JK, Harris J, Taylor SD, Palmer M. 2012. Characterization of daptomycin oligomerization with perylene excimer fluorescence: stoichiometric binding of phosphatidylglycerol triggers oligomer formation. Biochim Biophys Acta 1818:673–678.

70. Paradis-Bleau C, Kritikos G, Orlova K, Typas A, Bernhardt TG. 2014. A genome-wide screen for bacterial envelope biogenesis mutants identifies a novel factor involved in cell wall precursor metabolism. PLoS Genet 10:e1004056.

71. Djorić D, Kristich CJ. 2015. Oxidative stress enhances cephalosporin resistance of *Enterococcus faecalis* through activation of a two-component signaling system. Antimicrob Agents Chemother 59:159–169.

72. Sanders CC. 1988. Ciprofloxacin: *in vitro* activity, mechanism of action, and resistance. Rev Infect Dis 10:516–527.

73. Mager DL, Ximenez-Fyvie LA, Haffajee AD, Socransky SS. 2003. Distribution of selected bacterial species on intraoral surfaces: Microbiotas of intraoral surfaces. J Clin Periodontol 30:644–654.

74. Zehnder M, Guggenheim B. 2009. The mysterious appearance of enterococci in filled root canals. Int Endod J 42:277–287.

75. Cruz MR, Graham CE, Gagliano BC, Lorenz MC, Garsin DA. 2013. *Enterococcus faecalis* inhibits hyphal morphogenesis and virulence of *Candida albicans*. Infect Immun 81:189– 200.

76. Brown AO, Graham CE, Cruz MR, Singh KV, Murray BE, Lorenz MC, Garsin DA. 2019. Antifungal activity of the *Enterococcus faecalis* peptide EntV requires protease cleavage and disulfide bond formation. MBio 10:e01334–19.

77. Graham CE, Cruz MR, Garsin DA, Lorenz MC. 2017. *Enterococcus faecalis* bacteriocin EntV inhibits hyphal morphogenesis, biofilm formation, and virulence of *Candida albicans*. Proc Natl Acad Sci U S A 114:4507–4512.

78. Jurakova V, Farková V, Kucera J, Dadakova K, Zapletalova M, Paskova K, Reminek R, Glatz Z, Holla LI, Ruzicka F, Lochman J, Linhartova PB. 2023. Gene expression and metabolic activity of *Streptococcus mutans* during exposure to dietary carbohydrates glucose, sucrose, lactose, and xylitol. Mol Oral Microbiol 38:424–441.

79. Choi A, Dong K, Williams E, Pia L, Batagower J, Bending P, Shin I, Peters DI, Kaspar JR. 2024. Human saliva modifies growth, biofilm architecture, and competitive behaviors of oral streptococci. mSphere 9:e0077123.

80. Pultar F, Hansen ME, Wolfrum S, Böselt L, Fróis-Martins R, Bloch S, Kravina AG, Pehlivanoglu D, Schäffer C, LeibundGut-Landmann S, Riniker S, Carreira EM. 2021. Mutanobactin D from the human microbiome: Total synthesis, configurational assignment, and biological evaluation. J Am Chem Soc 143:10389–10402.

81. Hansen ME, Yasmin SO, Wolfrum S, Carreira EM. 2022. Total synthesis of mutanobactins A, B from the human microbiome: Macrocyclization and thiazepanone assembly in a single step. Angew Chem Int Ed Engl 61:e202203051.

82. Xie Z, Okinaga T, Qi F, Zhang Z, Merritt J. 2011. Cloning-independent and counterselectable markerless mutagenesis system in *Streptococcus mutans*. Appl Environ Microbiol 77:8025–8033.

83. Schindelin J, Arganda-Carreras I, Frise E, Kaynig V, Longair M, Pietzsch T, Preibisch S, Rueden C, Saalfeld S, Schmid B, Tinevez J-Y, White DJ, Hartenstein V, Eliceiri K, Tomancak P, Cardona A. 2012. Fiji: an open-source platform for biological-image analysis. Nat Methods 9:676–682.

84. Deatherage DE, Barrick JE. 2014. Identification of mutations in laboratory-evolved microbes from next-generation sequencing data using *breseq*. Methods Mol Biol 1151:165– 188.

85. Cisar JO, Kolenbrander PE, McIntire FC. 1979. Specificity of coaggregation reactions between human oral streptococci and strains of *Actinomyces viscosus* or *Actinomyces naeslundii*. Infect Immun 24:742–752.

86. Kilian M, Mikkelsen L, Henrichsen J. 1989. Taxonomic Study of Viridans Streptococci: Description of *Streptococcus gordonii* sp. nov. and Emended Descriptions of *Streptococcus sanguis* (White and Niven 1946), *Streptococcus oralis* (Bridge and Sneath 1982), and *Streptococcus mitis* (Andrewes and Horder 1906). Int J Syst Bacteriol 39:471–484.

87. Ajdić D, McShan WM, McLaughlin RE, Savić G, Chang J, Carson MB, Primeaux C, Tian R, Kenton S, Jia H, Lin S, Qian Y, Li S, Zhu H, Najar F, Lai H, White J, Roe BA, Ferretti JJ. 2002. Genome sequence of *Streptococcus mutans* UA159, a cariogenic dental pathogen. Proc Natl Acad Sci U S A 99:14434–14439.

88. Qi F, Chen P, Caufield PW. 2001. The group I strain of *Streptococcus mutans*, UA140, produces both the lantibiotic mutacin I and a nonlantibiotic bacteriocin, mutacin IV. Appl Environ Microbiol 67:15–21.

89. Markham JL, Knox KW, Wicken AJ, Hewett MJ. 1975. Formation of extracellular lipoteichoic acid by oral streptococci and lactobacilli. Infect Immun 12:378–386.

90. Sales MJ, Herbert WG, Du Y, Sandur AS, Stanley NM, Jensen PA. 2018. Complete genome sequences of *Streptococcus sobrinus* SL1 (ATCC 33478 = DSM 20742), NIDR 6715-7 (ATCC 27351), NIDR 6715-15 (ATCC 27352), and NCTC 10919 (ATCC 33402). Microbiol Resour Announc 7:e00804–18.

91. Dunny GM, Brown BL, Clewell DB. 1978. Induced cell aggregation and mating in *Streptococcus faecalis*: evidence for a bacterial sex pheromone. Proc Natl Acad Sci U S A 75:3479–3483.

92. Willett JLE, Ji MM, Dunny GM. 2019. Exploiting biofilm phenotypes for functional characterization of hypothetical genes in *Enterococcus faecalis*. NPJ Biofilms Microbiomes 5:23.

93. Sifri CD, Mylonakis E, Singh KV, Qin X, Garsin DA, Murray BE, Ausubel FM, Calderwood SB. 2002. Virulence Effect of *Enterococcus faecalis* Protease Genes and the Quorum-Sensing Locus *fsr* in *Caenorhabditis elegans* and Mice. Infection and Immunity 70:5647–5650.

94. Willett JLE, Robertson EB, Dunny GM. 2022. The phosphatase Bph and peptidyl-prolyl isomerase PrsA are required for gelatinase expression and activity in *Enterococcus faecalis*. J Bacteriol 204:e0012922.

95. Huycke MM, Spiegel CA, Gilmore MS. 1991. Bacteremia caused by hemolytic, high-level gentamicin-resistant *Enterococcus faecalis*. Antimicrob Agents Chemother 35:1626–1634.

96. Clewell DB, Tomich PK, Gawron-Burke MC, Franke AE, Yagi Y, An FY. 1982. Mapping of *Streptococcus faecalis* plasmids pAD1 and pAD2 and studies relating to transposition of Tn917. J Bacteriol 152:1220–1230.

97. Gilmore MS, Rauch M, Ramsey MM, Himes PR, Varahan S, Manson JM, Lebreton F, Hancock LE. 2015. Pheromone killing of multidrug-resistant *Enterococcus faecalis* V583 by native commensal strains. Proc Natl Acad Sci U S A 112:7273–7278.

98. Leuck A-M, Johnson JR, Dunny GM. 2014. A widely used *in vitro* biofilm assay has questionable clinical significance for enterococcal endocarditis. PLoS One 9:e107282.

99. Heaton MP, Handwerger S. 1995. Conjugative mobilization of a vancomycin resistance plasmid by a putative *Enterococcus faecium* sex pheromone response plasmid. Microb Drug Resist 1:177–183.

100. Kristich CJ, Manias DA, Dunny GM. 2005. Development of a method for markerless genetic exchange in *Enterococcus faecalis* and its use in construction of a *srtA* mutant. Appl Environ Microbiol 71:5837–5849.

101. Geldart K, Kaznessis YN. 2017. Characterization of Class IIa Bacteriocin Resistance in *Enterococcus faecium*. Antimicrob Agents Chemother 61:e02033–16.

102. Pollitt EJG, Szkuta PT, Burns N, Foster SJ. 2018. *Staphylococcus aureus* infection dynamics. PLoS Pathog 14:e1007112.

103. Kreiswirth BN, Löfdahl S, Betley MJ, O’Reilly M, Schlievert PM, Bergdoll MS, Novick RP. 1983. The toxic shock syndrome exotoxin structural gene is not detectably transmitted by a prophage. Nature 305:709–712.

104. Boles BR, Thoendel M, Roth AJ, Horswill AR. 2010. Identification of genes involved in polysaccharide-independent *Staphylococcus aureus* biofilm formation. PLoS One 5:e10146.

105. Wörmann ME, Reichmann NT, Malone CL, Horswill AR, Gründling A. 2011. Proteolytic cleavage inactivates the *Staphylococcus aureus* lipoteichoic acid synthase. J Bacteriol 193:5279–5291.

106. Bryan EM, Bae T, Kleerebezem M, Dunny GM. 2000. Improved vectors for nisin-controlled expression in gram-positive bacteria. Plasmid 44:183–190.

